# Structure and mechanism of microtubule stabilization and motor regulation by MAP9

**DOI:** 10.1101/2025.11.17.688911

**Authors:** Burak Cetin, Aryan Taheri, Mert Golcuk, Brigette Y. Monroy, Jonathan Fernandes, Kassandra M. Ori-McKenney, Mert Gur, Eva Nogales, Ahmet Yildiz

## Abstract

Microtubule-associated proteins (MAPs) regulate the organization of microtubules and control intracellular transport, but their individual contributions to microtubule dynamics and motor regulation remain poorly understood. Here, we identify MAP9 as a critical factor that stabilizes microtubules and facilitates neuronal morphogenesis. MAP9 knockdown abolishes the outgrowth of neurites, a phenotype not observed through the loss of other neuronal MAPs. Cryo-electron microscopy revealed that, unlike other MAPs that bind along protofilaments, MAP9 binds around the microtubule as a long alpha helix using five consecutive repeats. This unique binding mode enables MAP9 to staple adjacent protofilaments, thereby preventing microtubule depolymerization. We also showed that MAP9 selectively permits kinesin-3 and dynein motility while hindering kinesin-1 through interactions with a divergent loop-8 of the kinesin motor domain. Our results establish MAP9 as a key MAP required for neuronal growth and uncover how it differentially regulates intracellular transport driven by kinesin motors.

## Introduction

The microtubule (MT) cytoskeleton is the architectural backbone of neurons and supports long-range intracellular transport driven by kinesin and dynein. Structural MAPs such as tau, MAP2, MAP7, and doublecortin exhibit differential localization across neuronal compartments, creating functional domains such as the axon initial segment^1^. Traditionally, MAPs have been viewed as essential stabilizers of neuronal MTs, providing structural support to axons and dendrites and protecting MTs from depolymerization^2, 3^. However, recent studies have begun to challenge this long-standing view. Genetic depletion or knockout of individual MAPs, including tau, MAP1B, and even combinations of MAPs, often results in surprisingly mild or context-dependent phenotypes, without the widespread MT destabilization^4–7^. These observations led to the proposals that MAPs have overlapping functions in stabilizing MTs, such that the loss of one MAP can be compensated for by others^1^, or MAPs serve regulatory roles in MT organization rather than acting as generic stabilizers^8^. These ideas have not been carefully tested, and it remains unclear if other MAPs play indispensable, non-redundant roles in driving early axonal extensions.

MAP9, also known as Aster Associated Protein (ASAP), is a highly conserved MAP expressed in mitotic cells, cilia, and neurons^9–13^. MAP9 stabilizes and organizes the MT network in these cell types. Depletion of MAP9 causes defects in mitotic spindle organization, ultrastructural defects in axonemes, and embryonic lethality or severe malformations during neural development^9–13^. These findings hinted that MAP9 may play an essential, non-redundant role in MT organization across diverse developmental contexts. However, MAP9 is understudied compared to other MAPs, and the molecular basis for its contributions to MT regulation remains poorly understood.

MAP9 is known to differentially regulate kinesin motors involved in intracellular cargo transport. Unlike MAP7, which activates kinesin-1 motility^6, 14, 15^ and inhibits kinesin-3^16^, MAP9 has the opposite effect, inhibiting kinesin-1 while enabling kinesin-3 motility^16^. The mechanism by which MAPs distinguish between kinesin motors that are closely related is still emerging. Functional studies demonstrated that MAP7 selectively activates kinesin-1 by interacting with the coiled-coil stalk of the motor^6, 14–16^. However, MAP9 does not appear to strongly interact with either of these two kinesins, and it remains unclear how it can distinguish between different kinesin motors.

Here, we identify MAP9 as a critical regulator of early neuronal growth and intracellular transport, acting through a distinct MT-binding mode and motor-selectivity mechanism. Using cryo-electron microscopy (cryo-EM), we show that MAP9 binds MTs through a repeating motif that engages up to five adjacent protofilaments simultaneously, forming a lattice-stabilizing belt that suppresses MT catastrophe. In neurons, MAP9 knockdown abolishes neurite extension, highlighting its essential, non-redundant role in early neuronal morphogenesis. MAP9 does not overlap with kinesin on the MT, but it is positioned close to loop-8 of the kinesin motor domain. Insertion of loop-8 of kinesin-3 enabled kinesin-1 to walk uninhibited by MAP9, demonstrating that MAP9 selectively enables kinesin-3-driven transport by interacting with loop-8 of the kinesin motor domain. Together, these findings reveal MAP9 as a unique neuronal MAP that couples MT stabilization with selective motor regulation to promote early neuronal development.

## Results

### MAP9 binds across five protofilaments on the MT

Prior structural studies of MAP2, MAP4, MAP7, and tau had shown that these MAPs bind along a single protofilament^15, 17–19^. To determine if MAP9 also binds to the MT in this fashion, we expressed full-length (FL) human MAP9, which contains a disordered projection domain (PD) and an MT binding domain predicted to form a 200-residue-long α-helix (MTBD; **Fig. 1a, Supplementary Table 1**)^9^. Purified MAP9 is a monomer in solution and decorates MTs with a dissociation constant (K_D_) of 61 ± 10 nM (±SE; **Fig. 1b-d**). To determine the structure and MT binding mode of MAP9, we decorated MTs with excess FL MAP9 and obtained a cryo-EM reconstruction using the pseudo-helical symmetry of MTs (**Fig. 1e, Extended Data Figs. 1-2, Supplementary Table 2**). The reconstruction revealed that MAP9 binds to MTs through a single extended α-helix, consistent with the structure prediction of its MTBD. MAP9 binds across multiple protofilaments encircling the MT (**Fig. 1e**). Symmetry expansion of two laterally adjacent tubulins revealed how an 81-residue-long segment of α-helix within MAP9 MTBD binds to β- tubulin across the two neighboring tubulin dimers near the longitudinal interdimer interface (**Fig. 1f**).

**Fig. 1.**
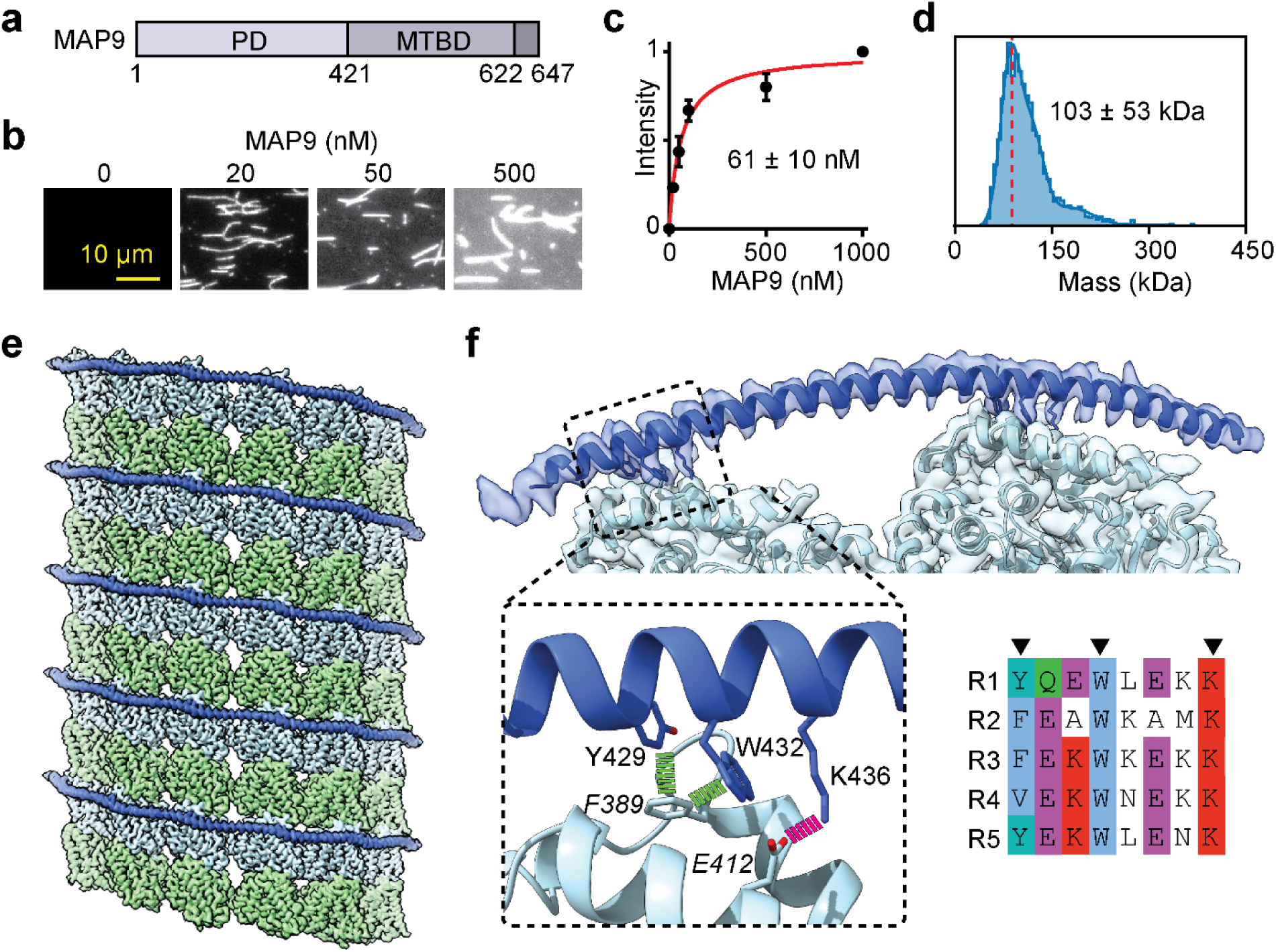
MAP9 bridges adjacent protofilaments of MTs. a, Domain schematic showing that MAP9 contains an N-terminal PD and a C-terminal MTBD. **b,** MT binding of fluorescently labeled MAP9 in BRB80 buffer (estimated ionic strength: 185 mM^42^). **c,** The fluorescence intensity (mean ± s.d.) of MAP9 was fitted to a noncooperative (hyperbolic) binding curve to calculate K_D_ (± SE; n = 30 MTs for all data points; two technical replicates). **d,** Mass photometry of purified GFP- MAP9 (mean ± s.d.). The red dashed line represents the expected monomer mass. **e,** Cryo-EM reconstruction of a MAP9-decorated MT obtained by imposing pseudo-helical symmetry. α- tubulin, β-tubulin, and MAP9 are shown in green, light blue, and royal blue, respectively. **f,** (Top) End-on view of the MAP9-MT cryo-EM reconstruction obtained by symmetry expansion using two adjacent protofilaments. (Bottom right) Alignment of the five MT binding pseudorepeats within MAP9 MTBD. (Bottom left) The fitted atomic model is shown as ribbons with all-atom representations of the side chains involved in hydrophobic (green) and ionic (magenta) interactions between the R1 pseudo-repeat of MAP9 (regular font) and β-tubulin (italic).

The interaction of MAP9 with β-tubulin involves a newly identified and well-conserved repeating sequence ФXXWXXXK, where Ф is mostly tyrosine or phenylalanine, W is tryptophan, K is lysine, and X is any amino acid (**Fig. 1f, Supplementary Fig. 1**). Ф and W form hydrophobic interactions with F389, and K forms a salt bridge with E412 of β-tubulin (**Fig. 1f**). MAP9 MTBD contains five of these pseudo-repeat sequences separated by 35-36-35-36 residue spacers, allowing them to reach across neighboring protofilaments and placing the pseudo-repeat in register with tubulin after 10 turns of the helix (**Fig. 2a**). Therefore, the 300 Å α-helix of MAP9 can bind up to five protofilaments. In our cryo-EM classifications, MAP9 does not bind α-tubulin, and its selectivity for β-tubulin likely arises from electrostatic differences at the binding interface, where β-tubulin Glu412 is replaced by α-tubulin Arg422 (**Extended Data Fig. 2d**). Lack of binding to α-tubulin indicates that MAP9 does not cross the seam of MTs (**Extended Data Fig. 2e**).

**Fig. 2.**
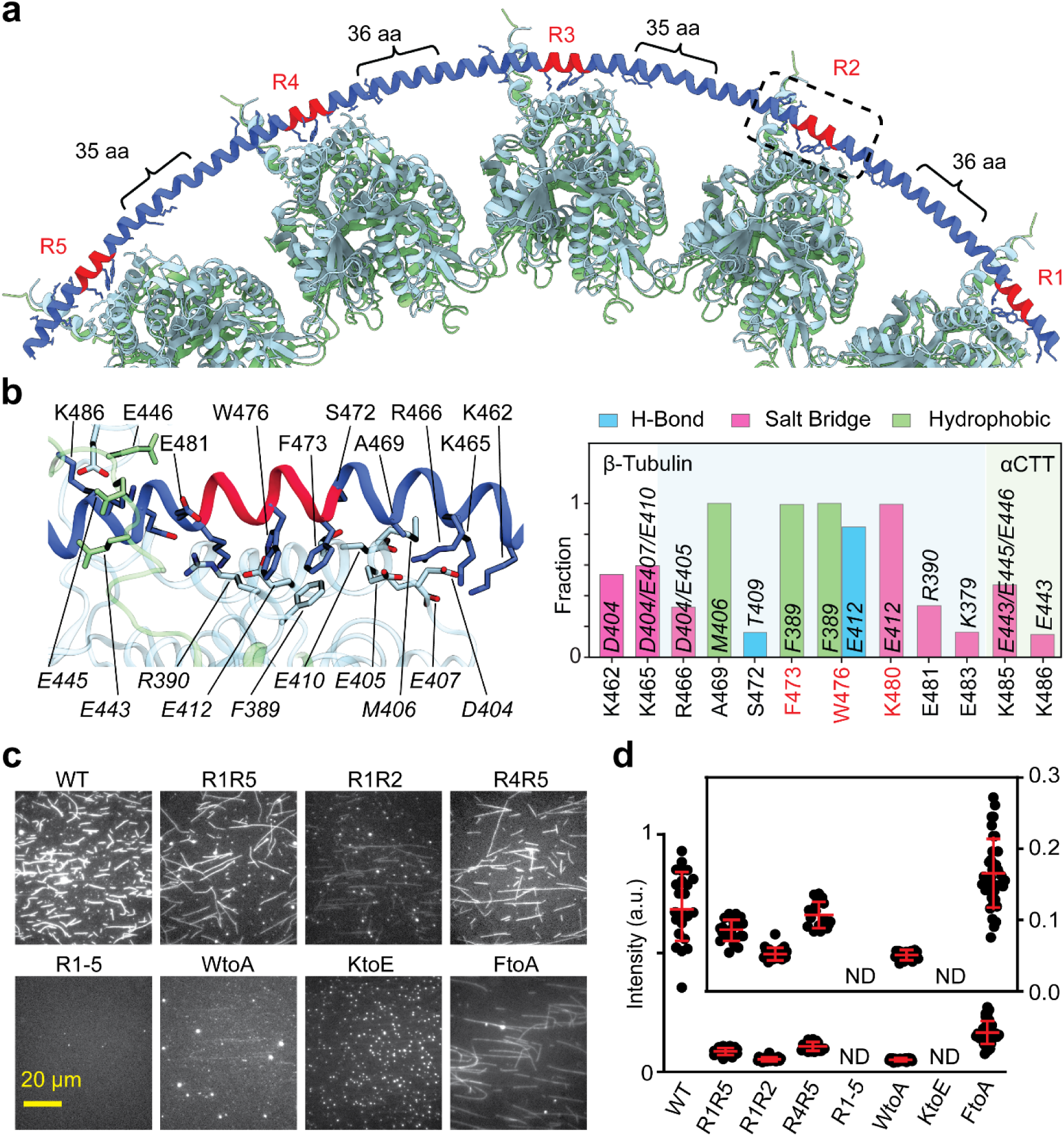
Pseudorepeats of MAP9 are critical for MT binding. a, Ribbon representation of the model of the MAP9 helix bound to five adjacent protofilaments through its pseudorepeats. **b,** (Left) A representative snapshot of MD simulations highlights the interactions between the R2 pseudorepeat of MAP9 (dark blue) and the tubulin dimer (α-tubulin in green and β-tubulin in blue). (Right) The fraction of specific intermolecular interactions formed between MAP9 and β-tubulin observed over the course of the MD simulations. **c,** MT decoration of 100 nM WT and mutant MAP9 constructs (see **Extended Data** Fig. 4a). **d,** The fluorescence intensity of WT and mutant forms of MAP9- MTBD on MTs (From left to right, n = 23, 22, 22, 36, 22, 22 MTs). The inset is the zoomed-in version of mutant MAP9 intensities. R indicates all three conserved residues were mutated in the specified pseudo-repeat (ND: not determined).

All-atom molecular dynamics (MD) simulations (1 µs in total length**, Supplementary Table 3**) confirmed that all five repeats of MAP9 bind identical residues in β-tubulin as they bridge neighboring protofilaments (**Fig. 2b, Supplementary Video 1**). In addition to the pairwise interactions that we identified from the cryo-EM reconstruction, simulations detected hydrogen bonding between conserved W and E412 of β-tubulin, hydrophobic interactions of MAP9 with M406 of β-tubulin, and transient electrostatic interactions with negatively charged residues in the α-tubulin’s C-terminal tail (CTT; **Fig. 2b, Extended Data Fig. 3**).

To understand how the identified repeats and pairwise interactions affect the MT binding of MAP9, we generated mutations in the binding repeats of a MAP9 construct containing only the MTBD (MAP9-MTBD, **Extended Data Fig. 4a**) and tested MT binding of wild-type (WT) and mutant MAP9-MTBD. Alanine substitutions or charge reversal of the conserved residues within the pseudo-repeats almost fully eliminated MT binding to 100 nM MAP9-MTBD (**Fig. 2c-d**). Mutating the three conserved residues in only two pseudo-repeats in varying combinations also substantially (∼80%) reduced MT binding (**Fig. 2d, Extended Data Fig. 4a-b**), demonstrating that all repeats contribute to MT binding of MAP9. In comparison, mutating the residues that contact αCTT did not affect MT binding (**Extended Data Fig. 4c-e**), likely because these interactions are relatively infrequent (21 ± 5% of the simulation time, **Extended Data Fig. 3**) and thus contribute less to the overall MT affinity of MAP9.

### MAP9 stabilizes dynamic MTs

The unique binding of MAP9 across protofilaments suggests a strong effect of this MAP on MT dynamics. To test this possibility, we monitored how MAP9 regulates MT polymerization and shrinkage in the presence of GTP and free tubulin in vitro and compared the effect of MAP9 to that of tau and MAP7. We observed that 300 nM MAP9 substantially promotes MT polymerization by doubling the MT growth rate (**Fig. 3a-b, Supplementary Video 2**). Unlike previous reports^20, 21^, 100 nM tau or 300 nM MAP7 did not substantially affect the MT growth rate but reduced the shrinkage rate of MTs. Remarkably, MAP9 increased the MT growth rate nearly two-fold, and slowed the shrinkage rate more than tau and MAP7 (**Fig. 3b**). Consistent with the bridging of neighboring protofilaments by this MAP, MAP9 nearly eliminated MT catastrophe, whereas MAP7 and tau had little to no effect on protecting MTs against catastrophes (**Fig. 3b**). These results contrast with earlier reports of the MT stabilization role of tau and MAP7^2, 22, 23^, and are more consistent with recent studies showing that tau does not significantly affect MT stabilization in neurons^24^.

**Fig. 3.**
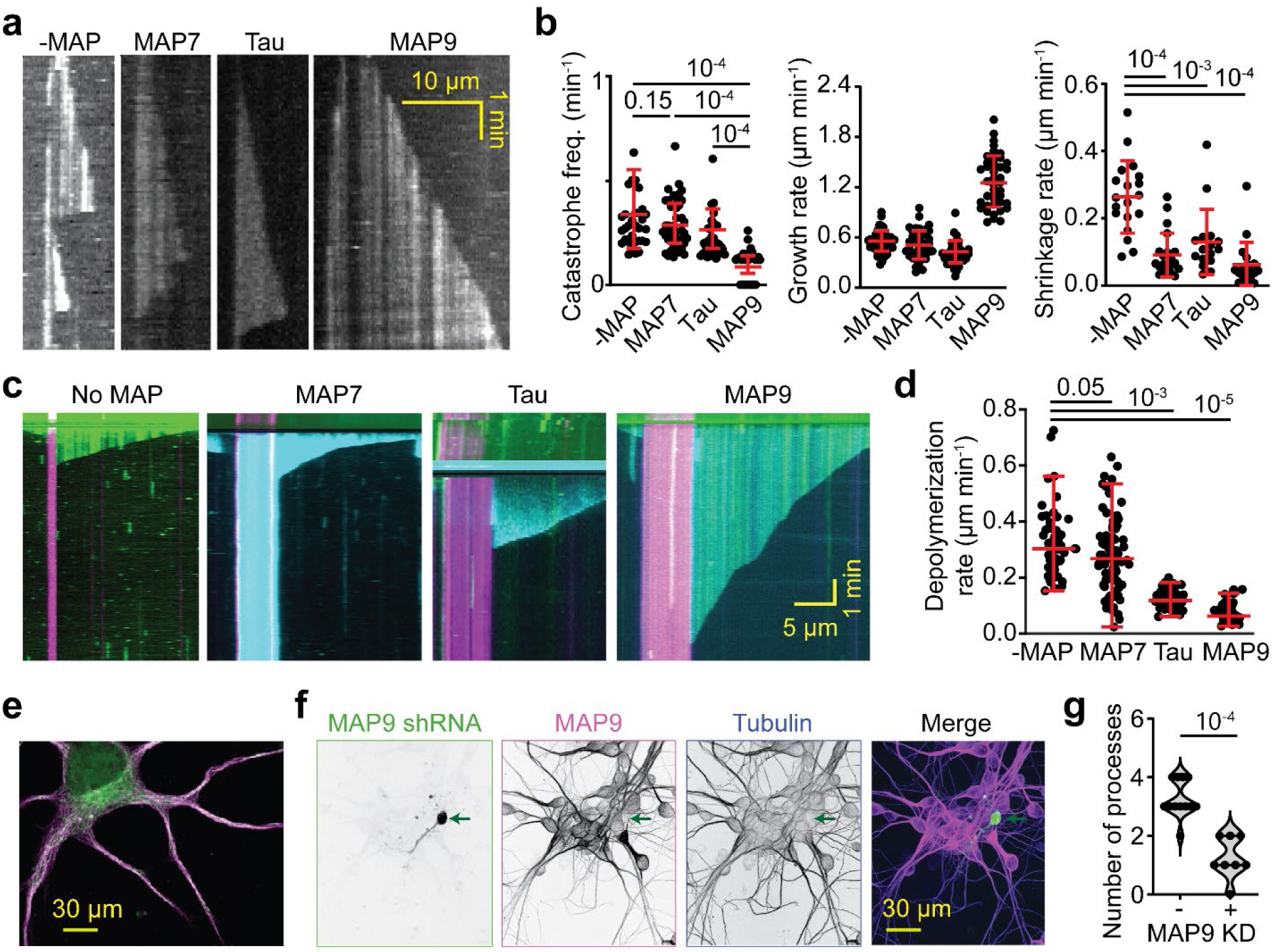
MAP9 promotes polymerization and suppresses the catastrophe of dynamic MTs. a, Kymographs of dynamic MTs in the presence or absence of 300 nM MAP7, tau, or MAP9. **b,** The catastrophe frequency, growth rate, and shrinkage rate of dynamic MTs in the presence and absence of MAPs (mean ± s.d.; from left to right, n =33, 41, 34, and 39 MTs for catastrophe; 56, 46, 43, and 42 MTs for growth rate; and 19, 21, 16, and 19 MTs for shrinkage rate; two technical replicates). **c,** Representative kymographs of MT depolymerization in the presence and absence of MAPs. MTs were polymerized from GMP-CPP seeds (magenta). MT extensions and MAPs are shown in green and cyan, respectively. **d,** Depolymerization rates of MTs after removing free tubulin and GTP in the flow chamber in the presence and absence of 300 nM MAP7, 300 nM MAP9, and 100 nM tau (from left to right, n = 57, 98, 75, 55 MTs, two technical replicates). **e,** Immunostaining of primary cultured neurons with antibodies against MAP9 (magenta) and α- tubulin (green). **f,** Confocal image of DIV4 neurons transfected with MAP9 shRNA. Arrows show that the MAP9 knockdown neuron (green) has only one neurite and is positive for tubulin (blue), but not for MAP9 (pink). **g,** The number of neurite processes extending from the cell body of DIV4 neurons in control and MAP9 shRNA-transfected neurons (n = 12 and 8 neurons, respectively, from 2 independent trials). In b and d, the centerline and whiskers represent mean and SD, respectively. In b and g, p-values were calculated from Welch’s two-tailed t-test.

To directly observe whether these MAPs protect MTs against depolymerization, we removed free tubulin and GTP but kept the MAPs in the assay chamber after growing dynamic MTs (**Fig. 3c**). In the absence of MAPs, MTs depolymerized at 0.3 µm s^-1^ (Ref. ^25^). MAP9 and tau slowed the depolymerization rate by 6-fold and 2.5-fold, respectively, whereas MAP7 failed to slow MT depolymerization (**Fig. 3d**). These results show that, among the MAPs we tested, MAP9 is most effective in stabilizing MTs and protecting them against depolymerization.

### MAP9 is essential for neurite growth

We next tested whether MAP9 is important for MT stability in neurons. Immunostaining in primary neuronal cultures revealed that MAP9 is distributed throughout neuronal extensions and is enriched in both axons and dendrites (**Fig. 3e, Extended Data Fig. 4f**), supporting its role in shaping polarized MT networks^16^. While total tubulin immunofluorescence produced a prominent, diffuse signal in the soma due to the large pool of soluble tubulin^26^, MAP9 staining was relatively weak in the cell body and strongly enriched along neurite shafts. This differential localization suggests that MAP9 may not bind soluble tubulin, but instead selectively decorate polymerized MTs (**Fig. 3e**).

To quantify MAP9 depletion in neurons, we performed immunofluorescence analysis in primary neuronal cultures and measured MAP9 signal normalized to tubulin intensity on a per-neuron basis, providing an internal control for cell-to-cell variability. MAP9 RNAi neurons exhibited a MAP9/tubulin ratio of 2.4 ± 0.7 compared to 4.9 ± 0.7 (mean ± s.d.) in control neurons, corresponding to an ∼51% reduction in MAP9 protein levels. Residual MAP9 signal was observed in RNAi-treated neurons, consistent with partial depletion achieved by RNAi rather than complete loss of MAP9 expression. Remarkably, the shRNA knockdown of MAP9 nearly fully eliminated neurite outgrowth, as evidenced by a large reduction in the number of extended neurites in shRNA-positive neurons compared to non-transfected or control neurons (**Fig. 3f-g, Extended Data Fig. 4g-h**). These results indicate that MAP9 is a key regulator of neuronal growth through its distinct MT-binding mode that robustly stabilizes MTs.

### MAP9 differentially regulates MT motors

We next investigated how MAP9 binding to MTs affects the motility of MT motors in vitro. We expressed a constitutively active human kinesin-1 KIF5B (K560, amino acids 1-560), human kinesin-2 KIF3A homodimer^27^, FL human kinesin-3 KIF1A, and mammalian dynein/dynactin/BicDR1 (DDR complex^28^; **Fig. 4a**). Motility assays revealed that MAP9 increases the frequency of kinesin-3 runs on MTs up to 2-fold at concentrations lower than K_D_ of MAP9^16^, but higher concentrations of MAP9 slightly inhibited the motility (**Fig. 4b-c, Extended Data Fig. 5a, Supplementary Video 3**). The inhibition constant (K_i_ = 310 nM) was substantially higher than the K_D_ of MAP9, suggesting that this inhibition occurs at near saturation of the MT surface by MAP9. Similar results were obtained for tail-truncated kinesin-3 (**Extended Data Fig. 5b**), indicating that MAP9 regulates kinesin-3 motility through its motor domain. MAP9 did not affect dynein motility at low concentrations but reduced its run frequency at higher concentrations (K_i_ = 158 nM; **Supplementary Video 4)**.

**Fig. 4.**
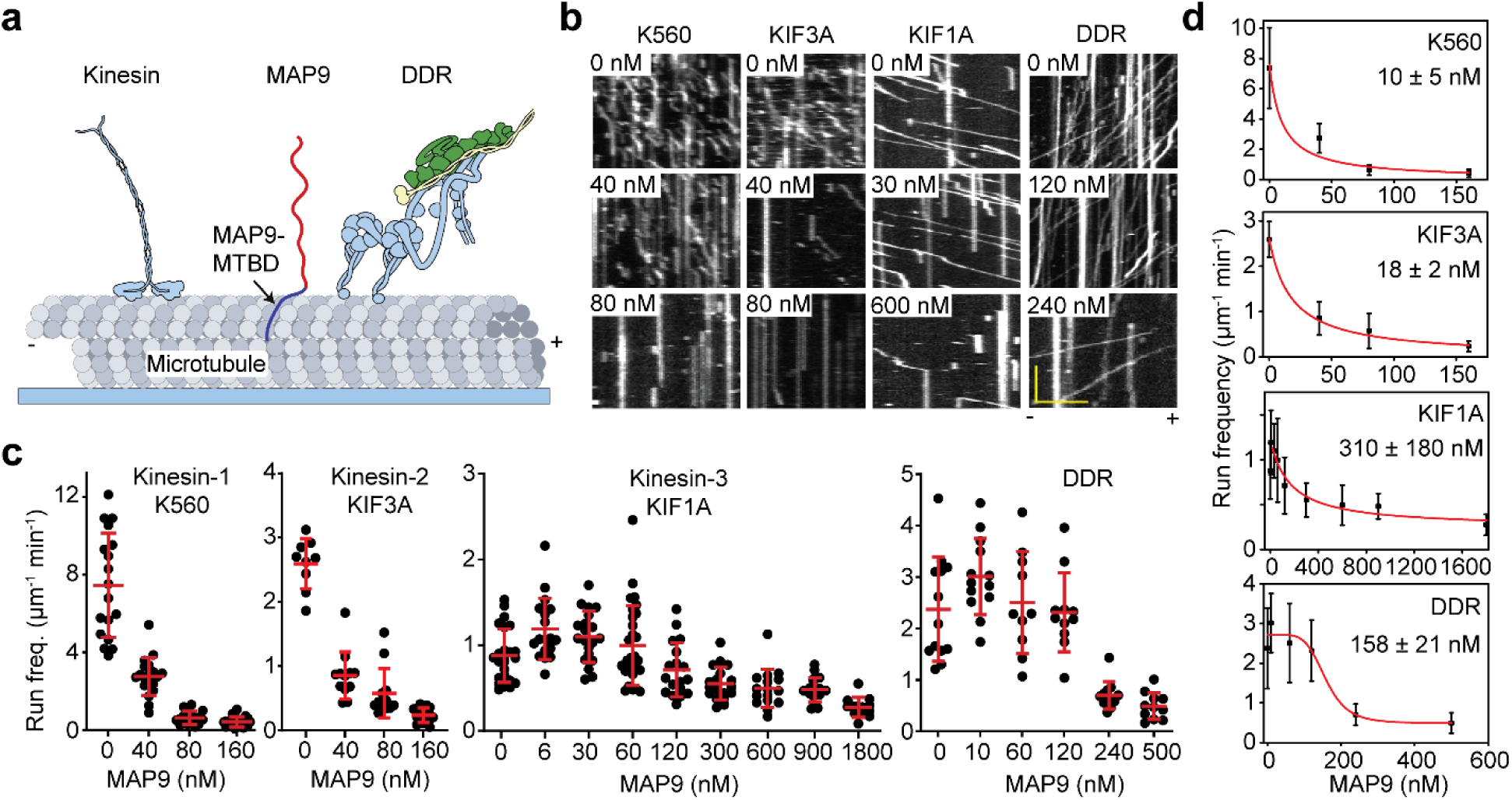
MAP9 differentially regulates kinesin and dynein motors. a, Schematic of single-molecule motility assays of kinesin and dynein (DDR) motors on MAP9-decorated MTs. The tail of KIF5B was truncated to generate a constitutively active motor (K560). **b,** Kymographs of K560, KIF3A, KIF1A, and DDR motility in the presence and absence of MAP9. **c,** Run frequency of K560, KIF3A, KIF1A, and DDR motors under different MAP9 concentrations (from left to right, n = 18, 19, 18, 22, 9, 12, 12, 15, 24, 19, 20, 29, 18, 19, 16, 18, 12, 13, 13, 10, 11, 12, 11 MTs). **d,** Run frequencies of K560, KIF3A, and KIF1A were fitted to a Langmuir isotherm, whereas the run frequency of DDR was fitted to the Hill Equation in their inhibitory form (red solid curves) to determine the inhibition constant (K_i_; ±SE).

Unlike kinesin-3 and dynein, kinesin-1 motility was strongly inhibited at concentrations lower than the K_D_ of MAP9 (K_i_ = 10 nM; **Supplementary Video 5)**, suggesting that approximately 20% saturation of the MT surface by MAP9 is sufficient to inhibit kinesin-1. MAP9 also substantially reduced the MT landing rate of kinesin-1 while not affecting the landing rate of kinesin-3 (**Extended Data Fig. 5c**), indicating that MAP9 primarily blocks MT binding of kinesin-1, rather than inhibiting the subsequent motility after its MT landing. Similar results were observed for kinesin-2 **(Fig. 4b-d, Extended Data Fig. 5c**), consistent with an increase in kinesin-2 velocity upon knocking out MAP9 in *C. elegans*^12^.

To understand why higher concentrations of MAP9 inhibit all motors, we performed motility experiments using MAP9-MTBD, which lacks the majority of the PD **(Fig. 5a, Extended Data Fig. 5d)**. MAP9-MTBD had similar MT binding affinity to full-length MAP9 **(Fig. 5b)**. MAP9- MTBD did not inhibit kinesin-3 or dynein motility at higher concentrations, while still inhibiting kinesin-1 motility as strongly as FL MAP9 (K_i_ = 14 nM; **Fig. 5c-d, Extended Data Fig. 5e, Supplementary Video 6)**. In addition, MAP9-MTBD resulted in an ∼80% increase in kinesin-3 run frequency. The activation constant (K_a_ = 60 nM) was similar to the K_D_ of MAP9 for MTs, suggesting that the increase in KIF1A run frequency by MAP9 is related to MT binding of MAP9. These results showed that the MTBD of MAP9 inhibits kinesin-1 while allowing kinesin-3 and dynein to walk on MTs. At near-saturation of MTs, the PD of MAP9 appears to reduce MT binding of motor proteins.

**Fig. 5.**
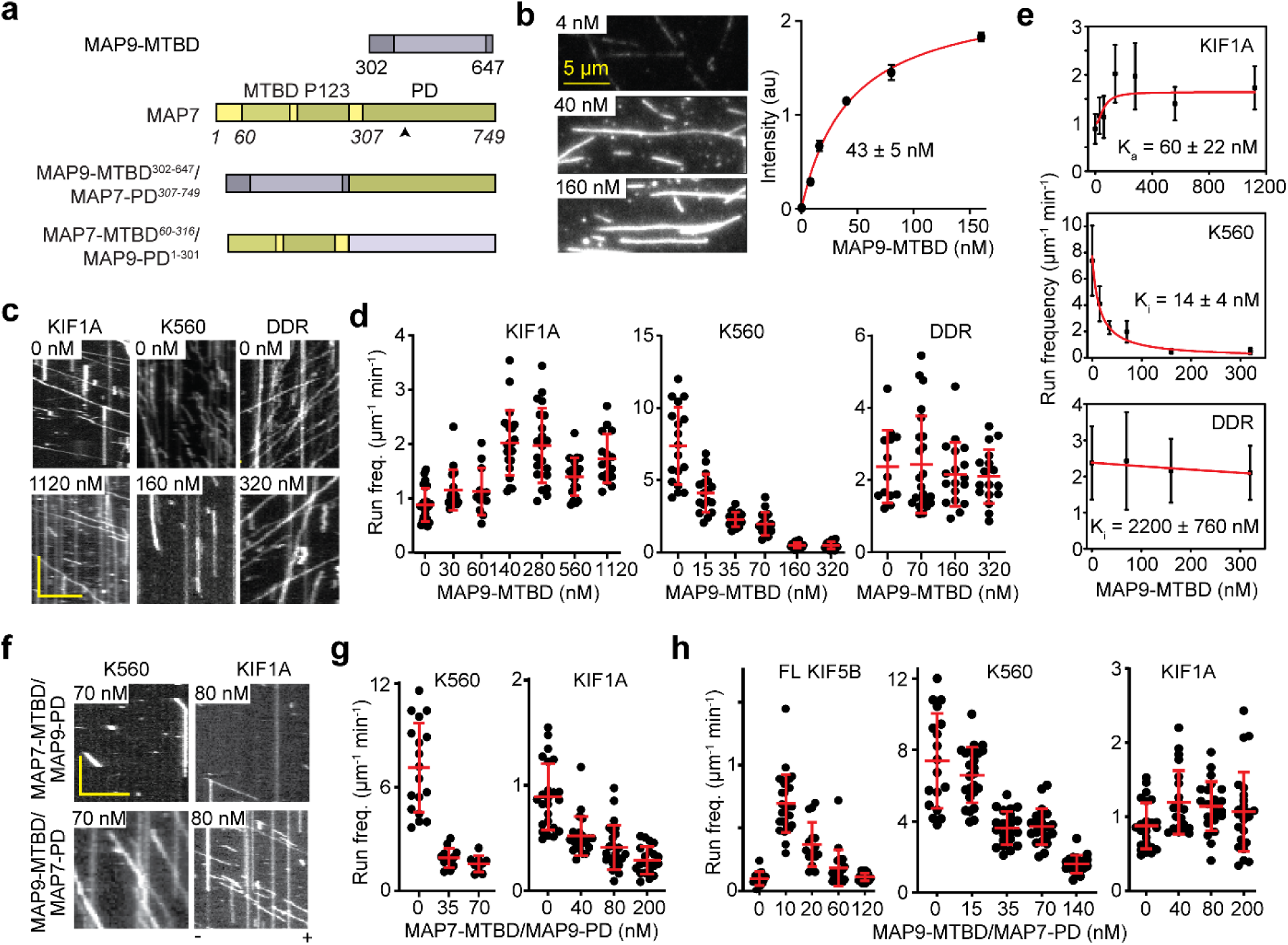
MAP9 differentially regulates kinesins through its MTBD. a, Schematic of WT and chimeric MAP7 and MAP9 constructs. The black arrow points to the KIF5B interaction site of MAP7 PD. **b,** (Left) MT decoration of fluorescently labeled MAP9-MTBD at different concentrations. (Right) The fluorescence intensity (mean ± SD) of MAP9-MTBD was fit to the Langmuir isotherm (solid curve) to estimate K_D_ (±se). From left to right, n = 28/29, 30/29, 36/32, 33/29, 30/29 MTs (two technical replicates). **c-d,** Kymographs and run frequency of KIF1A, K560, and DDR under different MAP9-MTBD concentrations (From left to right, n = 24, 17, 12, 22, 21, 19, 16, 18, 17, 15, 15, 19, 19, 13, 23, 18, 17 MTs). **e,** Run frequency of KIF1A was fitted to a Langmuir isotherm, whereas run frequencies of K560 and DDR were fitted to the Hill Equation in its inhibitory form (red solid curves) to determine the activation (K_a_) and inhibition constant (K_i_; ±SE). **f,** Kymographs (left) and run frequency (right) of KIF1A and K560 under different concentrations of chimeras MAP7-MTBD/MAP9-PD and MAP9-MTBD/MAP7-PD. **g,** Run frequency of KIF1A and K560 under different concentrations of chimeras MAP7-MTBD/MAP9- PD (from left to right, n = 18, 12, 9, 24, 22, 25, 21 MTs). **h,** Run frequency of FL KIF5B, K560, and KIF1A under different concentrations of MAP9-MTBD/MAP7-PD (from left to right, n = 15, 28, 15, 21, 24, 18, 25, 21, 21, 20, 24, 21, 23, 24 MTs). In c and f, the scale bars are 5 µm and 5 s on the *x* and *y* axes, respectively. In d, f, and g, the centerline and whiskers represent mean and SD, respectively.

To gain insight into the differential regulation of kinesin-1 and kinesin-3 by MAP9, we generated chimeric MAPs by fusing the MTBDs and PDs of MAP9 and MAP7^16^ **(Fig. 5a)**. In contrast to MAP9, MAP7 inhibits kinesin-3 while allowing kinesin-1 to walk on MTs **(Extended Data Fig. 6a-b)**^6, 14, 15^. The MTBD of MAP7 overlaps with the kinesin-binding site on tubulin, while its PD relieves kinesin-1 autoinhibition, acting as a tether to improve its landing on MTs^6, 14, 15^. A chimera of the MAP7 MTBD and MAP9 PD would therefore be expected to inhibit both motors, because this chimeric MAP overlaps with the motor binding site and lacks an activating tether for kinesin-1. In comparison, a chimera of MAP9 MTBD and MAP7 PD does not overlap with the kinesin binding site and contains an activating tether for kinesin-1. We anticipate this chimera to activate FL kinesin-1, but inhibit kinesin-1 motility at higher concentrations, similar to MAP7^15^. As expected, single-molecule motility assays revealed that MAP7 MTBD/MAP9 PD inhibited both kinesin-1 and kinesin-3 **(Fig. 5f-h, Extended Data Fig. 6c).** In comparison, a low concentration of MAP9 MTBD/MAP7 PD activated FL kinesin-1 motility, whereas it did not substantially affect constitutively active kinesin-1 (K560), similar to that observed for MAP7^15^. In addition, MAP9 MTBD/MAP7 PD did not substantially inhibit kinesin-3 motility, even at high concentrations, similar to MAP9 **(Fig. 5g,h, Extended Data Fig. 6c-e)**^16^. These results indicate that MAP9 differentially regulates kinesin-1 and kinesin-3 motility through its MTBD, not its PD.

### MAP9 distinguishes kinesins through their loop-8

It remained unclear how MAP9 could distinguish between kinesin-1 and -3, because these motors bind to MTs through a well-conserved motor domain within the kinesin family^29^. Docking the motor domains of these motors onto our map did not show an apparent steric clash with the MAP9 helix **(Extended Data Fig. 7).** To gain insight into how MAP9 regulates kinesins without an overlap on MTs, we obtained the cryo-EM structure of MTs co-decorated with MAP9 MTBD and the kinesin-3 motor domain and performed 3D focused classification of the symmetry-expanded map of tubulin dimers **(Fig. 6a-b, Extended Data Figs. 2 and 8)**. Unlike MAP7 MTBD, which precludes kinesin binding^15^, MAP9 binding allows kinesin-3 binding on the same tubulin dimer **(Fig. 6b)**.

**Fig. 6.**
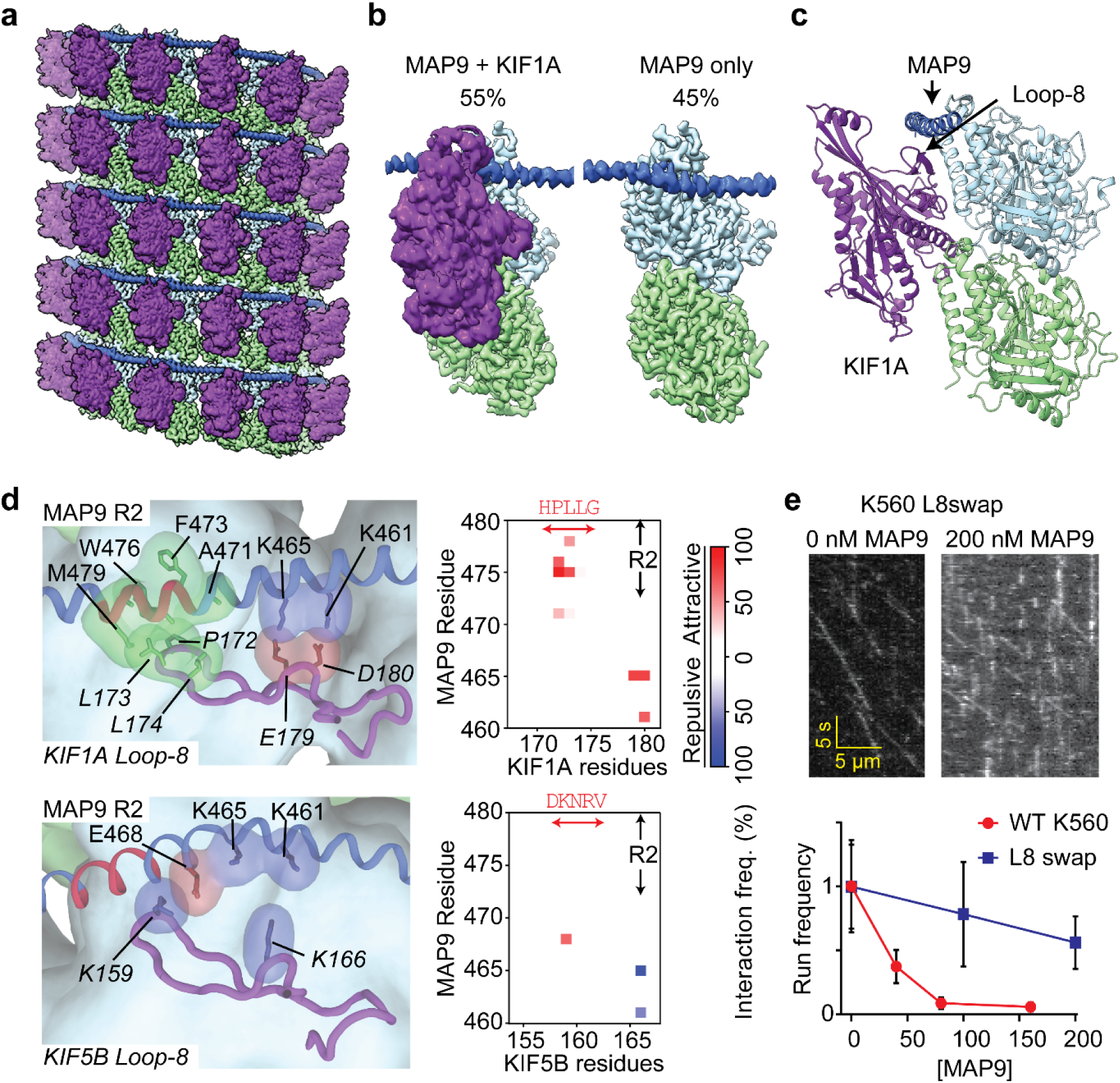
MAP9 distinguishes between kinesin-1 and -3 through loop8 of the kinesin motor domain. a, Cryo-EM reconstruction with pseudo-helical symmetry imposed for an MT co-decorated with MAP9 and the kinesin-3 motor domain. α-tubulin, β-tubulin, MAP9, and kinesin- 3 are shown in green, light blue, royal blue, and purple, respectively. **b,** Surface representation and percentage of the two classes resulting from symmetry expansion and 3D classification on a single tubulin dimer. **c,** Atomic model from the cryo-EM reconstruction of MAP9 and kinesin-3 bound to the same tubulin. **d,** (Left) Interactions between R2 of MAP9 and loop-8 of kinesin-3 (top) and kinesin-1 (bottom) in MD simulations. Hydrophobic, negative, and positive side chains are highlighted in green, red, and blue, respectively. (Right) The frequencies of attractive and repulsive interactions between kinesin and MAP residues are color-coded based on interaction frequency. The red arrow highlights the loop-8 residues used to generate the chimeric K560 L8swap motor. **e,** Kymographs (top) and the normalized run frequency (bottom) of the chimeric K560 L8swap in the presence and absence of MAP9 (±SD; from left to right, n = 15, 21, 22 MTs). The run frequency of WT K560 is shown for comparison.

The model built into the cryo-EM map showed that the MAP9 helix is positioned near loop-8 of the kinesin-3 motor domain **(Fig. 6c)**. Loop-8 of kinesin-3 consists of hydrophobic residues, whereas the equivalent region of kinesin-1 consists of positively charged residues **(Extended Data Fig. 7b-c)**. To test the possibility that differences in loop-8 between kinesin-1 and kinesin-3 are critical for their selective regulation by MAP9, we ran MD simulations of kinesin-1 and -3 motor domains interacting with each repeat of MAP9-MTBD on the MT (9.6 µs total simulation time; **Extended Data Figs. 9 and 10**). The presence of MAP9 did not substantially affect the interaction of kinesin-3 loop-8 with the MT, but slightly reduced the frequency of salt bridges formed between kinesin-1 loop-8 and β-tubulin (**Extended Data Figs. 9b and 10b**). However, we identified attractive and repulsive interactions between MAP9 and the two kinesins by identifying residue pairs that exhibited positively or negatively correlated motions at the interaction interface^30^. In kinesin-3, loop-8 formed favorable hydrophobic and ionic interactions with MAP9 **(Fig. 6d, Extended Data Fig. 9, Supplementary Video 7)**. In comparison, kinesin-1 loop-8 did not form hydrophobic interactions and exhibited increased negative correlations due to electrostatic repulsion from similarly charged residues in MAP9 **(Fig. 6d, Extended Data Fig. 10, Supplementary Video 8)**. To test these observations, we substituted five residues of loop-8 of kinesin-1 with those of kinesin-3 (K560 L8swap; **Fig. 6e**). This modification substantially reduced the inhibition of kinesin-1 by MAP9 when compared to the WT motor under the same MAP9 concentrations (**Fig. 6e, Extended Data Fig. 7d**), in agreement with our model that MAP9 differentially regulates kinesin-1 and -3 via the differences between their divergent loop-8 regions.

## Discussion

Our findings establish MAP9 as a structurally and functionally distinct MAP that not only stabilizes the MT cytoskeleton through a novel binding architecture but also encodes specificity in intracellular transport. While tau and MAP7 bind MTs along protofilaments^15, 18^, MAP9 adopts a unique circumferential binding mode, spanning up to five adjacent protofilaments. In our cryo-EM classifications, MAP9 only binds β-tubulin, likely because residues in β-tubulin that interact with the MAP9 pseudo-repeat are not conserved in α-tubulin. In particular, the β-tubulin E412 residue that forms a salt bridge with the conserved lysine in the MAP9 pseudo-repeat is replaced by α- tubulin R422. As a result, MAP9 cannot cross the seam while maintaining the same binding configuration observed across non-seam lateral interfaces, because such a configuration would require MAP9 to engage α-tubulin at the seam. Consistent with our results, a recent cryo-electron tomography study of the ciliary transition zone identified MAP9 as a B-tubule-specific MAP that decorates five protofilaments through contacts with β-tubulin^31^. These findings confirm the MAP9 binding footprint in situ and implicate MAP9 in both anterograde transport regulation and structural support of the cilium^31^.

The unique MT binding mode of MAP9 effectively “staples” the protofilaments together, dramatically promoting MT growth and suppressing MT catastrophe compared to tau and MAP7. Previous studies suggested that lateral contacts between protofilaments play a more significant role than longitudinal contacts in resisting depolymerization^32–34^. Our results provide direct evidence supporting this model. The ability of MAP9 to bind across protofilaments likely antagonizes protofilament peeling^32, 34^, thereby protecting MTs against depolymerization more effectively than those MAPs that stabilize longitudinal contacts within single protofilaments^15, 18^.

Consistent with this mechanistic difference, we show that depletion of MAP9 results in complete arrest of neurite outgrowth. In comparison, knockdown of tau or MAP7 induces axonal branching and dendritic phenotypes in mice, but these neurons were able to develop and extend both dendrites and axons without major perturbations to the overall morphology^4–7^. The contrast suggests that not all MAPs contribute equally to lattice stability in vivo and highlights MAP9 as an essential and nonredundant factor required for sustaining the structural framework that supports neurite extension. More broadly, these results challenge the long-held view that MAPs act as functionally interchangeable stabilizers^35^ and instead point to a division of labor in which the specific binding geometry of individual MAPs dictates their ability to preserve axonal and dendritic MTs. These insights position MAP9 as a key regulator of neuronal architecture and a promising target for future studies into the mechanisms of neuronal development. The physiological role of MAP9 in neurons has not been studied in detail, and further studies employing higher-efficiency depletion strategies or genetic loss-of-function approaches will be necessary to fully dissect the specific contributions of MAP9 to neuronal development and maintenance.

Our results also demonstrate how MAP9 exerts selective control over kinesin-1 and kinesin-3- driven transport without overlapping with their MT footprints. Previous functional assays attributed this selectivity to the presence of a tandem repeat of lysine residues within loop-12 of the kinesin-3 motor domain^16^, but the underlying mechanism remained unclear. Our cryo-EM studies revealed that the MAP9 helix does not directly overlap with the binding sites of dynein and kinesin motors at the intradimer interface of α- and β-tubulin. Instead, MAP9 is positioned at the plus-end-directed side of kinesin, facing towards loop-8 of the kinesin motor domain. Loop-8 exhibits sequence divergence among kinesins and has been identified as a determinant of MT binding and processivity of kinesin-3^36, 37^. Simulations and functional assays revealed that MAP9 forms attractive interactions with the loop-8 of kinesin-3, whereas its interactions with the loop-8 of kinesin-1 are repulsive, establishing the mechanistic basis of how MAP9 distinguishes between these motors despite their similarities. Substituting these residues in kinesin-1 with the corresponding kinesin-3 residues prevents inhibition by MAP9, showing that MAP9 inhibits kinesin-1 motility by recognizing its loop-8 sequence.

Differential regulation of the motors by MAP9 may also influence bidirectional transport. For cargos transported by dynein-dynactin and kinesin-1, MAP9 is expected to permit dynein motility while inhibiting kinesin-1, thereby favoring retrograde transport. In contrast, MAP9 is expected to have little to no effect on cargos transported by dynein and kinesin-3. Thus, MAP9 decoration of MTs may selectively inhibit bidirectional cargos that use kinesin-1 as the plus-end-directed motor, while allowing kinesin-3-driven cargos to move largely unimpeded along MAP9-decorated MTs.

Our work introduces a new regulatory principle for how MAPs recognize divergent sequences of kinesin motors to gate their transport. Sequence divergence in these loops contributes to differences in autoregulation^38^ and force generation^39^ of kinesin motors, as well as to their class-specific functions^29^, such as MT depolymerization^40, 41^. Our findings add a new layer to this regulatory code, showing that loop divergence can also serve as a recognition module for kinesin motors by MAPs.

## Acknowledgments

We thank R. McKenney for constructs, M. Aslan, J. Slivka, Y. Zhao and J. Fernandes for assistance with protein purification, J. Weng for assistance with fluorescence image acquisition and processing, J. Peukes for microscopy and computational support, M. Petersen and S. Marqusee for CD spectra, J. Al-Bassam for helpful discussions regarding MAP9 sequence motifs, the Cal-Cryo facility at UC Berkeley for EM imaging, and Indiana Jetstream2 and NCSA Delta for MD simulations. This work was supported by grants from the National Institute of General Medical Sciences (GM136414 (A.Y.), GM127018 (E.N.), GM133688 (K.M.O.M.)) and the National Science Foundation (MCB-1954449 and MCB-2534447, A.Y.; DGE-2146752, A.T.), Advanced Cyberinfrastructure Coordination Ecosystem (ACCESS, BIO240144, M.Gur). E.N. is a Howard Hughes Medical Institute Investigator.

## Author contributions

B.C., A.T., E.N., and A.Y. conceived the project and analyzed the data. B.C. purified the proteins and performed single-molecule experiments. A.T. performed cryo-EM sample preparation, data collection, and analysis. B.Y.M. and K.M.O.M. performed studies in cultured neurons. M. Gur designed and supervised MD simulations, and M. Golcuk performed MD simulations and analysis. B.C., A.T., M. Golcuk, K.M.O.M., M. Gur, E.N., and A.Y. wrote the manuscript.

## Methods

### Plasmids and Construct Design

The plasmid containing the FL KIF1A sequence with a C-terminal mScarlet fusion in a pOmniBac backbone was received from R. McKenney, UC Davis^43^. This tag was replaced with a yBBr tag using Gibson assembly for organic dye labeling. The DNA sequence that expresses the mouse kinesin-2 KIF3A motor domain and neck linker, dimerized through the kinesin-1 coiled-coil, was obtained from AddGene (Plasmid #129768). This sequence was cloned into a pET28(a) + vector with a C-terminal StrepII and yBBr tag for purification and organic dye labeling, using NdeI and XhoI restriction sites. Human KIF1A (1-393) fused to an artificial dimerization leucine zipper tag, ybbR labeling tag, and Twin-Strep purification tag and cloned into a pET28(a)+ vector using Gibson assembly. *E.coli* expression vectors for WT and mutant MAP9-MTBD constructs with C-terminal GFP and StrepII tags, chimeric MAP7/MAP9 constructs with N-terminal GFP and C-terminal StrepII tags, and monomeric KIF1A (residues 1-393) with an N-terminal mTurquoise2 and C-terminal StrepII tag were ordered from Twist Biosciences. Briefly, the open reading frame was cloned into the pET28a(+) vector using NdeI and XhoI restriction sites, which introduces a cleavable His-tag into the N-terminus of the expressed protein. The K560 L8swap construct with a C-terminal mScarlet fusion was cloned into the pET29b vector using the same restriction sites. The constructs used in this study are listed in **Supplementary Table 1**.

### Protein Expression and Purification

The plasmid for expression of FL human KIF1A with the pOmniBac backbone was transformed into DH10Bac competent cells, followed by plating onto Bacmid plates with BluoGal at 37 °C for 2 days under antibiotic selection. A white colony was selected and grown in LB medium overnight. Bacmid plasmids were isolated and transfected into adherent SF9 cells using Lipofectamine. The transfected SF9 cells were incubated at 27 °C for 3 days to grow the initial virus (P1). 2 mL of the P1 virus was added to 50 mL of suspended SF9 cell culture and incubated at 27 °C in a shaking incubator for 3 days. The P2 virus was collected by centrifuging at 4,000 *g* for 10 min and stored at 4 °C in the dark for several months. The amplified P2 virus was then used to infect a 2 L suspension SF9 culture, and the protein was expressed for 72 h in serum-containing media. The cells were harvested via centrifugation at 4,000 *g*, and the resulting pellet was snap-frozen in liquid nitrogen. The cell pellet was stored at -80 °C. To purify the protein, the cell pellet was thawed and resuspended in SFP buffer (50 mM HEPES-KOH, pH 7.4, 300 mM KCl, 10 mM MgCl_2_, 10% glycerol). The resuspended cells were lysed by a dounce homogenizer. The lysate was cleared via centrifuging at 150,000 *g* for 45 min in a Ti70 rotor (Beckman Coulter). The clarified lysate was then applied to 1 mL of Strep-Tactin XT resin equilibrated in the lysis buffer using gravity flow with disposable columns. The resin containing the Strep-tag protein was then washed with 20 column volumes of SFP buffer. The FL KIF1A was then fluorescently labeled with LD655-CoA dye (Lumidyne) using SFP synthase and SFP buffer. Briefly, a 2 mL solution containing 50 nmol dye and 10 µM SFP synthase in SFP buffer was passed through the column, which was then capped at both ends and incubated overnight at 4 °C. The resin was then washed with 10 column volumes of SFP buffer to wash away the free dye. The protein was then eluted with SFP buffer containing 50 mM d-biotin. Labeled aliquots of FL KIF1A were flash-frozen and stored at -80 °C.

Human KIF1A-LZ (residues 1-393) and mouse kinesin-2 KIF3A homodimer were expressed in *E. coli* BL21 (DE3) cells. Briefly, a 2L culture was grown in TB medium to an OD_600_ of 0.5, and protein expression was induced by adding 0.5 mM IPTG at 16 °C for 18 h. These proteins were purified and labeled identically as FL KIF1A, except the cells were lysed via tip sonication after harvesting.

Human MAP9 fused to GFP and a Twin-Strep tag (a gift from R. McKenney, UC Davis) and MAP9-MTBD were expressed in Rosetta (DE3) cells. Briefly, a 2 L culture was grown in TB medium to an OD_600_ of 0.5, and protein expression was induced by adding 0.5 mM IPTG at 16 °C overnight. The protein was purified via the Strep-II tag without overnight labeling and proceeded directly to elution after washing the resin with the lysis buffer. For structural studies, the eluate was then diluted threefold using SFP buffer and applied to a heparin HP column (Cytiva). The column was washed with an SFP buffer containing 100 mM KCl, followed by stepwise elution with an SFP buffer under increasing KCl concentration from 100 to 500 mM. The eluate was concentrated and further purified via gel filtration to remove other contaminants on a Superdex 200 column.

Pig-brain dynactin and the cargo adaptor mouse BicDR1 were purified as described previously^28^. The mutant of human dynein that cannot form the autoinhibited phi conformation (Phi mutant) was purified and fluorescently labeled as previously described^44^. The LD655-labeled single-use frozen aliquots were stored at -80 °C and used after thawing the same day. C-terminally His-tagged human KIF5B (K560) was purified as described previously^45^.

### Sample Preparation for Light Microscopy

Plain glass coverslips were cleaned with water, acetone, and water by sonication for 10 min, followed by sonication in a bath sonicator for 40 min in 1 M KOH. The coverslips were then rinsed with ddH_2_O, incubated in 3-Aminopropyltriethoxysilane in acetate and methanol for 10 min while sonicating for 1 min between successive steps, further cleaned with methanol, and finally air-dried. 30 µl of 25% biotin-PEG-succinimidyl valerate in a NaHCO_3_ buffer (pH 7.4) was sandwiched between two coverslips, followed by incubation at 4 °C overnight. The coverslips were then cleaned with water, air-dried, vacuum-sealed, and stored at -20 °C. Flow chambers were built by sandwiching a double-sided tape with a PEG-coated coverslip and a glass slide.

### Light Microscopy

The fluorescent imaging was performed with a custom-built multicolor objective-type TIRF microscope equipped with a Nikon Ti-E microscope body, a 100X magnification, 1.49 N.A. apochromatic oil-immersion objective (Nikon) with a Perfect Focus System. The fluorescence signal was detected using an electron-multiplying charge-coupled device camera (Andor, iXon EM+, 512 × 512 pixels). The effective camera pixel size after magnification was 160 nm. Alexa488/GFP/mNeonGreen, LD555, and LD655 probes were excited using 488 nm, 561 nm, and 633 nm laser beams (Coherent) coupled to a single-mode fiber (Oz Optics), and their emission was filtered using 525/40, 585/40, and 697/75 bandpass filters (Semrock), respectively. The microscope was controlled using Micromanager 1.4.

The flow chambers were incubated with a 5 mg mL^−1^ streptavidin solution for 3 min and washed with BRB80 buffer (80 mM PIPES pH 6.8, 1 mM EGTA, 2 mM MgCl_2_). The ionic strength of this buffer is 185 mM^42^. Assays were performed in the absence of additional salt. Biotinylated MTs were then added to the flow chamber and incubated for 3 min. The optimal concentration of biotinylated MTs was determined empirically. The chamber was then washed with BRB80 buffer supplemented with 1 mg mL^-1^ casein and 0.5% pluronic acid. Kinesins, motors, and MAP proteins were diluted in BRB80 supplemented with 1 mg mL^-1^ casein, 0.5% pluronic acid, and 2 mM ATP. Compared to KIF1A-LZ, a higher motor concentration (40X) of FL KIF1A was flown into the chamber to dimerize and activate the motor. Movies were recorded for ∼2-4 min at 200 ms per frame. Motors and ATP were not added for experiments measuring the fluorescence intensity of MAPs on MTs.

For DDR motility, 200 nM LD655-labeled phi-mutant dynein was incubated with 200 nM pig brain dynactin and 350 nM mouse BicDR1 in BRB80 buffer supplemented with 1 mg mL^-1^ casein, 0.5% pluronic acid, and 2 mM ATP. The complex was incubated for 15 min at room temperature and diluted 25-fold. The diluted dynein was added to the flow chamber and imaged on the microscope for 10 mins in the presence or absence of MAP9.

### Dynamic MT imaging

Biotinylated GMPCPP-labeled seeds were generated, as described previously^46^. Pig brain tubulin (30 mg mL^-1^) was mixed with 2% Cy5-labeled tubulin and centrifuged at 4 °C at 100,000 rpm using a TLA 120.1 rotor (Beckman). The supernatant was kept on ice and used for MT polymerization in a flow chamber. Flow chambers with a biotin-PEG coverslip were preincubated with 1 mg mL^-1^ streptavidin for 3 min and washed with BRB80 to remove unbound streptavidin. Biotin-labeled GMPCPP seeds were immobilized on the coverslip for 3 min, and the chamber was washed with BRB80 buffer containing 1 mg mL^-1^ casein and 0.5% pluronic acid. After immobilizing the seeds, the pig brain tubulin with a mixture of labeled and unlabeled tubulin was diluted 10-fold in BRB80 buffer supplemented with 4 mM GTP, 0.2% methylcellulose, 1 mg mL^-1^ casein, 0.5% pluronic acid, 0.1 mg mL^−1^ glucose oxidase, 0.02 mg mL^−1^ catalase, 0.8% D-glucose in the presence and absence of 300 nM MAP9. The mixture was flowed into the chamber and imaged immediately on the TIRF microscope at 22 °C. Movies were recorded for 100 frames with 5 s between frames for 8 min.

### Fluorescence Image Analysis

For the fluorescence intensity of MTs, images of MTs with MAP9 or MAP7 bound were imported into ImageJ, and the average fluorescence intensity per length of an MT was measured using the measure tool. The data were plotted in Prism, and the fluorescence intensity values were fitted to the Langmuir binding isotherm (non-cooperative) to determine the apparent K_D_.

Single-molecule motility of kinesins and dyneins was recorded for 250-1,000 frames per imaging area. A modified version of FIESTA (YFIESTA) was used to generate MT tracks and kymographs from recorded movies. YFIESTA was used to detect single processive trajectories in kymographs. Nonmotile, short (less than three pixels of movement), or diffusive trajectories (appearing as random forward and backward motion along the MT) were excluded from the analysis. In case of overlap between the trajectories in a kymograph, original TIF movies were manually analyzed to separate the movement of individual motors from each other.

The run frequency was calculated by observing the number of processive motors on each MT divided by the length of the MT and time. The run length and velocity were calculated by manually scoring the beginning and the end of single trajectories in the kymographs. In the presence of full-length MAP9, the run frequency (RF) of K560, KIF3A, and KIF1A in different MAP9 concentrations ([MAP9]) was fitted to a Langmuir isotherm in its inhibitory form (*RF*([*MAP*9]) = 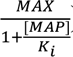), whereas the run frequency of DDR was fitted to the Hill Equation in its inhibitory form (*RF*([*MAP*9]) = *MIN* + 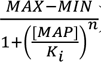), where *MAX* and *MIN* are the maximum and minimum run frequencies, *n* is the Hill coefficient, and *K_i_* is the inhibition constant. In the presence of MAP9- MTBD, the run frequency of KIF1A was fitted to a Langmuir isotherm (*RF*([*MAP*9]) = *MIN* +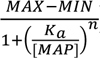) to calculate the activation constant (*K_a_*), whereas run frequencies of K560 and DDR were fitted to the Hill Equation in its inhibitory form to estimate *K_i_*. Fits were performed in Origin.

### Circular Dichroism (CD)

CD spectra were collected at room temperature on a JASCO J-1500 spectropolarimeter using proteins at 1–5.5 μM in 10 mM sodium phosphate (pH 7.4) and 150 mM KCl. Measurements were performed in quartz cuvettes with 1 cm or 1 mm path lengths. Spectra were collected in triplicate and normalized to the signal at 220 nm prior to comparison. Data acquired at HT voltages exceeding 600 V were excluded from the analysis due to noise.

### Neuronal Culture Isolation

For neuronal cultures, hippocampi were dissected out from E16.5–18.5 mouse embryonic brains, and hippocampal neurons were isolated using the Worthington Papain Dissociation System (Worthington Biochemical Corporation) according to the manufacturer’s protocol. Primary neurons were plated at a density of 500,000 cells per glass coverslip coated with 0.05 mg/mL Poly-D-Lysine (molecular weight 70,000–150,000, Millipore-Sigma) and cultured in Neurobasal media supplemented with glucose, GlutaMAX (ThermoFisher), B27, and penicillin/streptomycin (P/S). The media was changed every two days until neurons were fixed at DIV4. A high plating density was used because MAP9 knockdown led to substantial neuronal loss, resulting in relatively few MAP9 shRNA-transfected neurons.

### RNA Interference

To knock down MAP9, we used the GIPZ Lentiviral shRNA glycerol set (Horizon Discovery/Dharmacon), which targets the mouse MAP9 gene. The GIPZ shRNA sequences have been validated by the manufacturer for efficient and specific knockdown of MAP9 expression in mammalian cells. The set consisted of six different shRNA constructs (in pGIPZ vectors) with the following clone IDs and mature antisense sequences: RMM4431-200312303, Clone ID: V2LMM_124251, Antisense: TCCGTTCAATTCTTTCTTG RMM4431-200314601, Clone ID: V2LMM_124254, Antisense: TCTCTTTGAGGTATTCTAG RMM4431-200348107, Clone ID: V2LMM_240138, Antisense: TTCTTTGAAGGAAGATGAG RMM4431-200353088, Clone ID: V2LMM_211213, Antisense: ATGTGATCAGCCTTGTCTG RMM4431-200368801, Clone ID: V3LMM_418357, Antisense: CTAACTAGACTTTCATCTG RMM4431-200365512, Clone ID: V3LMM_418356, Antisense: TTGATAGCAAGGAATCCCA

All six constructs were combined in equal ratios (1:1) to generate a pooled shRNA mixture. This mixture was transfected into primary hippocampal neurons using Lipofectamine 2000 (ThermoFisher) according to the manufacturer’s instructions. Transfection efficiency was assessed using the internal GFP reporter cassette included in the GIPZ vector. Neurons were fixed at DIV4 and analyzed for MAP9 expression and morphological changes.

Quantification of MAP9 levels was performed on a per-neuron basis. For each neuron, regions of interest were defined around the cell body and along a proximal neurite, and mean fluorescence intensities for MAP9 and tubulin were measured using Fiji. MAP9 intensity was normalized to tubulin intensity for each neuron to account for cell-to-cell variability and staining efficiency. The resulting MAP9/tubulin ratio was used as a relative measure of MAP9 protein abundance.

### Immunocytochemistry

The cultures were fixed in 4% paraformaldehyde for 20 min at room temperature, washed several times with phosphate-buffered saline (PBS), permeabilized with 0.3% Triton X-100 in PBS (PBS- TX), and blocked with 5% BSA in PBS for 1 h at room temperature. Cultures were then incubated overnight at 4 °C with primary antibodies: rabbit anti-MAP9 (1:500, Invitrogen PA5-58145) and mouse anti-alpha tubulin (1:1000, Sigma Clone DM1A, T9026). The next day, cultures were incubated with secondary antibodies (1:1000) for 1 h at room temperature: Cy3 donkey anti-rabbit, Cy5 donkey anti-mouse, or Cy3 goat anti-chicken. Following several PBS washes, coverslips were mounted using VectaShield mounting medium (Vector Laboratories). Imaging was performed using a Leica SPE laser scanning confocal microscope with a 60x or 100x oil immersion objective. The total number of neuronal processes was quantified in GFP-positive MAP9 shRNA-transfected neurons and compared to control neurons.

### Cryo-EM Sample Preparation

Lyophilized porcine brain tubulin (Cytoskeleton) was resuspended to 10 mg mL^-1^ in BRB80 buffer supplemented with 10% (v/v) glycerol, 1 mM GTP, and 1 mM DTT. 10 μL of the tubulin solution was polymerized at 37 °C for 15 min. 1 μL of 2 mM taxol was added to the tubulin solution and incubated at 37 °C for 10 min. 1 μL of 2 mM taxol was added again, and the solution was incubated for 30 min. MTs were pelleted by centrifugation at 37 °C and 17,000 rcf for 20 min. The supernatant was discarded, and the pelleted MTs were resuspended in resuspension buffer (BRB80 buffer supplemented with 250 μM taxol). An aliquot of the resuspended MTs was depolymerized by CaCl_2_ to measure the concentration of polymerized tubulin. The remaining MT solution was diluted to 1.5 μM in dilution buffer (BRB80 buffer supplemented with 100 μM taxol). Immediately before sample preparation, all MT-binding proteins were desalted to BRB80 using Zeba Spin columns (Pierce).

To prepare MT-bound MAP9 samples, 2 μL of 1.5 μM taxol-MTs were incubated on a glow-discharged holey carbon cryo-EM grid (QuantiFoil, Cu 300 R 1.2/1.3) for 30 s, manually blotted with Whatman filter paper, and 2 μL of 7 μM MAP9 was added to the grid. The grid was transferred to a Vitrobot (ThermoFisher) set at 25 °C and 80% humidity, plunge-frozen in liquid ethane with a blot force of 5 pN and a blot time of 6 s, and transferred to liquid nitrogen.

MTs were decorated with MAP9-MTBD plus KIF1A (1-393) following a similar procedure. 2 μL of 1.5 μM taxol-MTs were incubated on a glow-discharged holey carbon cryo-EM grid for 30 s, manually blotted as before, and incubated with 2 μL of 6 μM MAP9-MTBD, manually blotted, then incubated with 6 μM KIF1A (1-393) in 2 mM AMP-PNP. The grid was plunge-frozen in the Vitrobot under identical conditions as the previous sample and transferred to liquid nitrogen.

### Cryo-EM Data Collection

Cryo-EM data for MT-bound MAP9 and MT-bound MAP9-MTBD and KIF1A (1-393) were collected on an Arctica microscope (ThermoFisher), operated at 200 kV with a K3 direct electron detector (Gatan). Images were acquired at 36,000x nominal magnification at 1.14 Å/pixel. All data were acquired in the super-resolution mode with a dose rate of ∼7.2 electrons/pixel/second and an exposure time of ∼9 s, dose-fractionated into 50 frames. All data were collected using the SerialEM software package^47^.

### Cryo-EM Image Processin**<u>g</u>**

For the MT-MAP9 dataset, the movie stacks were imported to CryoSPARC^48^ and motion-corrected (**Supplementary Table 2**). The CTF parameters were estimated with the patch CTF job and manually curated to remove bad micrographs. We followed the pipeline used in our previous studies, which deals with the pseudo-helical symmetry of MTs and uneven binding of multiple MAPs^15, 46^. Briefly, particles were automatically picked with the filament tracer, initially without a reference, and later using 2D templates as input^49^. The segment separation was set to 82 Å, the length of an αβ tubulin dimer. Particle images were initially extracted with a box size of 512 pixels and Fourier-cropped to 256 pixels. Four rounds of 2D classification were performed, and classes showing clear density for the MT were selected. MTs containing 13 or 14 protofilaments were separated by using initial models of 13- and 14-protofilament MTs as references in heterogeneous refinement. In both cryo-EM datasets, the majority of particles were classified as 14-protofilament MTs, so this class was used for downstream processing. 14-protofilament MT particles were subjected to helical refinement with an initial rise estimate of 82.5 Å and a twist of 0°. Particles were then subjected to local refinement using a cylindrical mask enclosing the MT. To obtain a reconstruction that accounts for the MT seam, we used a seam search routine that accounts for the fact that helical symmetry is broken at a site of heterologous lateral contacts between tubulins using Frealign^50^, with custom scripts that determine the seam position on a per-particle basis^51^. For this purpose, we converted the CryoSPARC alignment file from the last local refinement, first to the STAR format using the csparc2star script from the PyEM suite (10.5281/zenodo.3576630), and then to PAR Frealign format using a custom Python script^46^. Upon completion of the seam search protocol, the particles were imported back to CryoSPARC with the seam-corrected alignments and re-extracted without Fourier cropping (box size: 512 pixels). The imported particles were subjected to local refinement, and CTF refinement was performed to estimate the per-particle CTF. Another local refinement was run on this particle set to produce the final C1 reconstruction. The symmetry search job was used to determine the rise and the twist of the map, and these parameters were input to exploit the pseudo-symmetry of the MT. A local refinement was performed on the newly expanded particle stack with a cylindrical mask around the MT to yield the final symmetrized reconstruction of the entire MT. A final local refinement was performed with a smaller mask around the good protofilament (i.e., the protofilament opposite to the seam) and the protofilament adjacent to it, producing an approximately 3Å resolution map of MAP9-bound MTs.

The MT-MAP9-MTBD-KIF1A (1-393) dataset was processed using the same protocol to produce a consensus reconstruction at 2.9Å resolution. To identify particle distributions associated with MAP9, KIF1A, or both MAP9 and KIF1A co-bound to MTs, 3D classification using four classes was performed in CryoSPARC with a more constrained mask covering only one tubulin dimer, one KIF1A monomer, and the MAP9 region covering a single tubulin dimer. The 3D classification of four classes yielded two classes with MAP9 bound to a tubulin dimer without a KIF1A monomer and two classes with MAP9 and a KIF1A monomer co-bound to a tubulin dimer. The particles in similar classes were merged into the two described states, each comprising approximately half of the data: MAP9 alone on MTs, and both the KIF1A monomer and MAP9 co-bound to MTs.

### Model Building and Refinement

The KIF1A-MAP9-MT structure was modeled by first fitting the deposited model of the two-head-bound KIF1A AMPPNP-MT structure (PDB ID: 8UTO^39^). Once the model was adjusted to fit the structure, it was morphed using ISOLDE^52^. The MAP9 R1-R2 segment was modeled into the density map by first identifying the MAP9 residues involved in MT binding across the first two pseudorepeats. The main identifying feature was the clear tryptophan residue protruding into the β-tubulin density. The AlphaFold2-predicted MAP9 model^53^ was then tailored to include only the start of the MTBD and an extension matching the length of the helix identified in the density map. This tailored model was initially fitted into the map and subsequently morphed using ISOLDE. The identified MT-binding pseudorepeats fit well within the density map. Next, the model was combined with the adjusted KIF1A-MT model, and the combined structure was subjected to real-space refinement in Phenix^54^. The MAP9 R1-R2 (with a 36 a.a. spacing between pseudorepeats) and the MAP9 R2-R3 (with a 35 a.a. spacing between pseudorepeats)-MT model were generated by fitting tubulins (PDB: 6DPV) and tailoring the AlphaFold2 MAP9 model to extend the helix containing the residues identified as the first and second or the second and third MT-binding pseudorepeats, respectively. This extended helix was fitted and morphed into the density map using ISOLDE. Finally, both the MAP9 R1-R2-MT and MAP9 R2-R3-MT models underwent real-space refinement in Phenix^54^.

### MD Simulations

An atomic model of the MAP9 MT-binding domain (MTBD; residues A421–H624) bound to five (α-β-α) protofilaments, including the tubulin CTTs, was built based on the cryo-EM reconstruction. Additional models were generated in which the MTs were co-decorated with MAP9-MTBD and KIF5B or KIF1A motor domains (**Supplementary Table 3a**). The kinesin-MT systems were built using one tubulin dimer. All systems were solvated in a TIP3P water box, ensuring at least 30 Å of water padding in each dimension. Ions were added to neutralize the systems and set final concentrations to 150 mM KCl and 1 mM MgCl_2_. The kinesin-MAP9-MT systems contained approximately 1.5 M atoms, while the kinesin-MT systems contained 150 K atoms. MD simulations were performed using NAMD3^55^ and the CHARMM36m^56^ all-atom additive protein force field. The simulations used a 2-fs time step at 310 K and 1 atm. Long-range electrostatics were treated with the particle-mesh Ewald method, and a 12 Å cutoff was applied for van der Waals interactions.

To equilibrate the system, the protein was first held fixed during 10,000 steps of minimization, followed by 1 ns of equilibration. The protein constraints were then released, and the system underwent an additional 10,000 steps of minimization. Subsequently, a 5-ns equilibration was performed while restraining the C_α_ atoms with a harmonic potential of 1 kcal mol^−1^Å^−2^. Following these two minimization-equilibration phases, the production run was initiated. To account for the missing structural elements of MAP9 and the MT, C_α_ atoms of select tubulin residues (**Supplementary Table 3b**) were restrained with a harmonic potential of 1 kcal mol^−1^Å^−2^. Eight simulation runs of 125 ns in length each were performed for the MAP9-MT complex, eight runs of 700 ns each were conducted for the kinesin-3-bound MAP9-MT complex, and eight runs of 500 ns each were carried out for the kinesin-1-bound MAP9-MT complex. After the initial equilibration period, both the kinesin-1-MAP9-MT and kinesin-3-MAP9-MT systems reached stable root-mean-squared displacement (RMSD) plateaus, typically within 3-5 Å (**Supplementary Figure 2**), with no major conformational rearrangement or disruption of the MAP9-MT or kinesin- MAP9-MT interfaces. Production trajectories were concatenated for ensemble-level interaction-frequency and correlation analyses. To assess robustness across independently initiated simulations, trajectory-specific contact correlation matrices were compared to verify that the key MAP9-kinesin and kinesin-MT interaction features recur across simulations (**Supplementary Figures 3-4**). Additional simulations were performed in the absence of MAP9 to quantify how the presence of MAP9 affects kinesin-MT interactions (**Supplementary Figure 5**).

Correlations between the amino acids of MAP9 and kinesin were evaluated using a three-step method^30, 57^. First, we constructed an inter-protein covariance matrix C, defined as C = 〈(R − 〈R〉)(R − 〈R〉)^T^〉, where **R** represents the configuration vector containing only the C_α_ coordinates for a given MD simulation conformation, and ⟨**R**⟩ denotes the trajectory-averaged configuration vector. Next, we filtered residue pairs based on their interatomic distances and finally focused on the correlated residues in close contact to pinpoint the specific residue interactions responsible for these observed correlations. Salt bridges were identified as interactions between basic nitrogen and acidic oxygen atoms within 6 Å^58^. Hydrogen bonds were defined as interactions where the donor–acceptor distance was ≤3.5 Å and the donor–hydrogen–acceptor angle was ≤30°^59^. Hydrophobic interactions were defined as contacts between side-chain carbon atoms within an 8 Å cutoff distance. Repulsive correlations were identified when distances between pairs of either basic nitrogen or acidic oxygen atoms were within 12 Å.

### Statistical analysis

*P* values were determined using statistical tests in Prism. Two-sided two-sample t-tests were performed to compare continuous datasets, assuming that data are approximately normally distributed. Each result was based on a minimum of two independent experiments. The number of replicates and data points (n) and the specific statistical methods used are provided in the figure legends. Representative data from independently repeated experiments are shown.

## Data Availability

A reporting summary for this article is available as a Supplementary Information file. The main data supporting the findings of this study are available within the article and its Extended Data Figures. Protocols supporting the findings of this study are described in the Methods section. The constructs that express WT and mutant versions of MAP9 will be deposited at AddGene. Raw microscopy data will be made available by the corresponding authors upon request. The coordinates for MAP9 R1 and R2 bound to tubulin, MAP9 R2 and R3 bound to tubulin, and MAP9 and the KIF1A motor domain bound to tubulin are available at the Protein Data Bank (PDB) with accession codes 9ZP6, 9ZP7, and 9ZP5, respectively. All cryo-EM maps are available at the EMDB with accession codes EMD-74518 and EMD-74517.

## Code availability

The custom code used to analyze experimental data is uploaded to Zenodo (doi: 10.5281/zenodo.18041601). Scripts for cryo-EM image processing are available from E.N. upon request.

## Extended Data Figures

**Extended Data Fig. 1.**
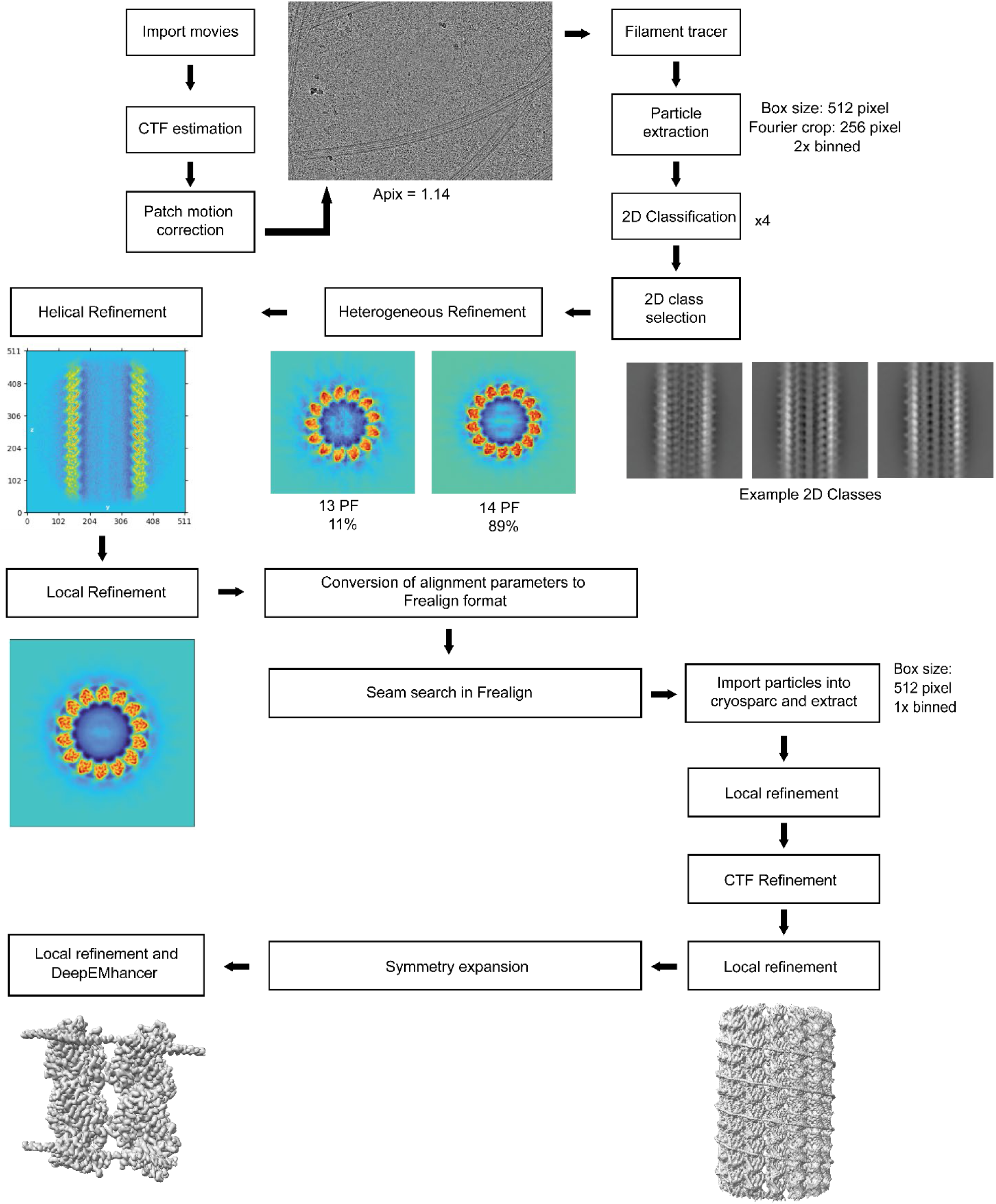
Cryo-EM data processing pipeline used to determine the structure of MAP9 bound to MTs.

**Extended Data Fig. 2.**
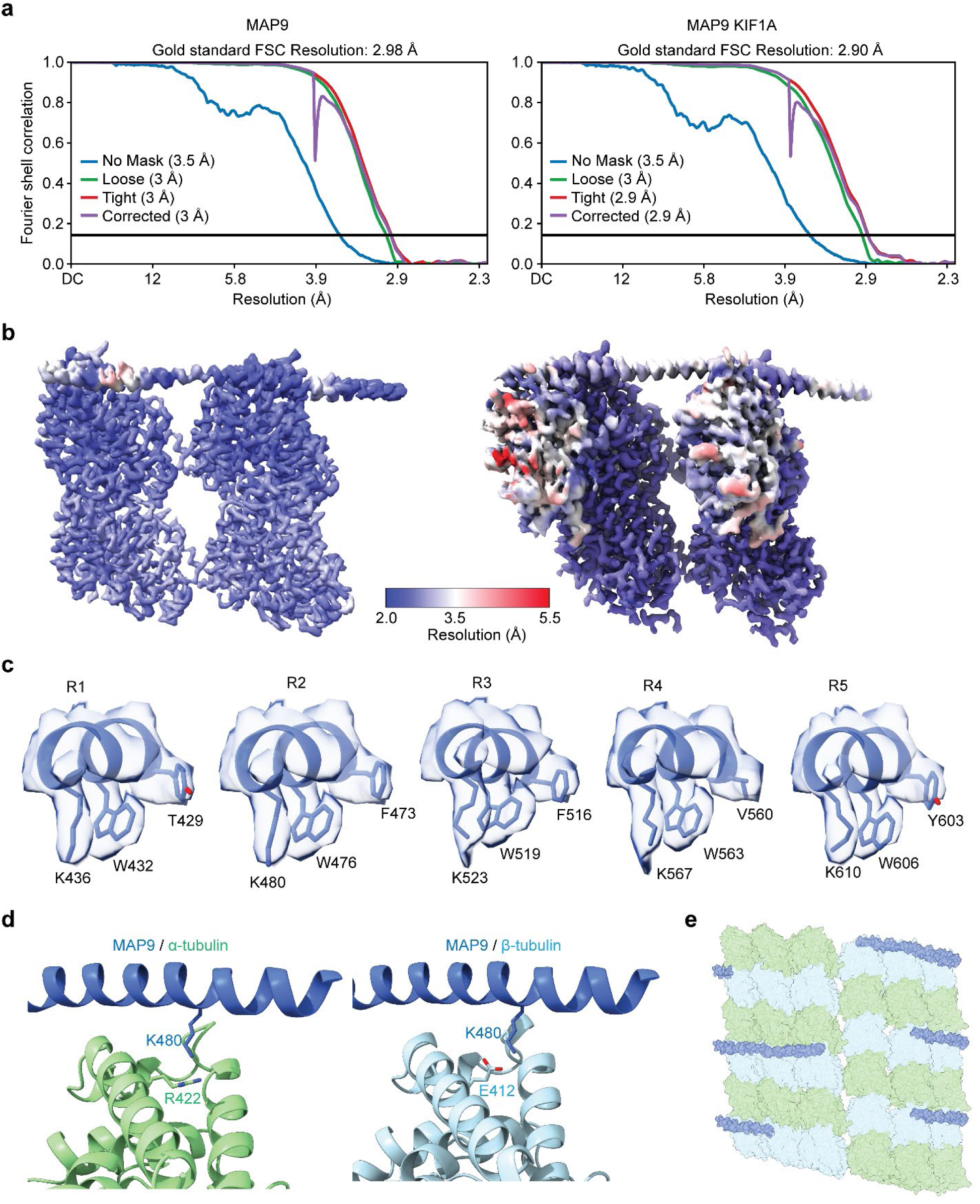
Resolution of the cryo-EM maps. **a**, Fourier shell correlation (FSC) plots of symmetrized reconstructions of MAP9 bound to MTs (left) and MAP9 and the KIF1A motor domain bound to MTs (right). **b,** Resolution maps of the symmetrized reconstruction of MAP9 bound to MTs (left) and MAP9 and the KIF1A motor domain bound to MTs (right). **c,** Fit of the five MT-binding domains of MAP9 into the cryo-EM density. The views of repeats 1, 2, and 3 are built atomic models within the cryo-EM density map. Repeats 4 and 5 are fitted atomic models into the density map. **d,** The fit of the α-tubulin model onto the density of β-tubulin shows a charge reversal in a key MAP9 interaction site with β-tubulin. **e,** Cartoon schematic of MAP9 stochastic binding to β-tubulins across protofilaments, but not crossing the seam.

**Extended Data Fig. 3.**
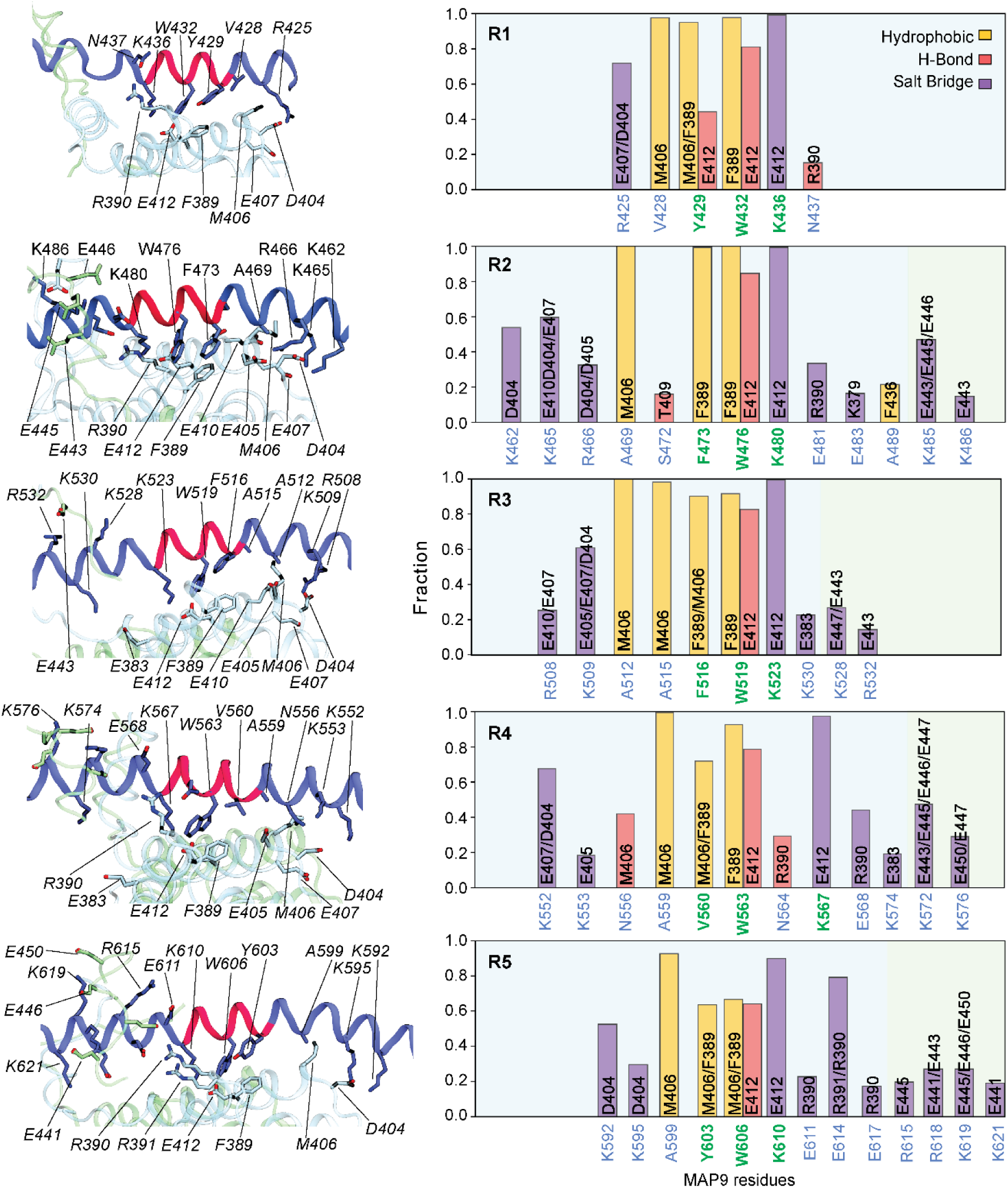
Interactions between MAP9 and tubulin observed in all-atom MD simulations. (Left) Representative structural snapshots highlight the interactions between MAP9 (dark blue) and the tubulin dimer (α-tubulin in green and β-tubulin in blue). The protein backbones are depicted as new cartoons, while interacting side chains are shown in licorice representation. (Right) The fraction of specific interactions formed between MAP9 and β-tubulin observed over the course of the MD simulations. The positively charged residues on MAP9 also transiently interacted with the glutamic acid side chains of the β-tubulin tail (light yellow; E441, E443, E445, E446, E447, E450). Conserved MAP9 residues in pseudo-repeats are highlighted in green.

**Extended Data Fig. 4.**
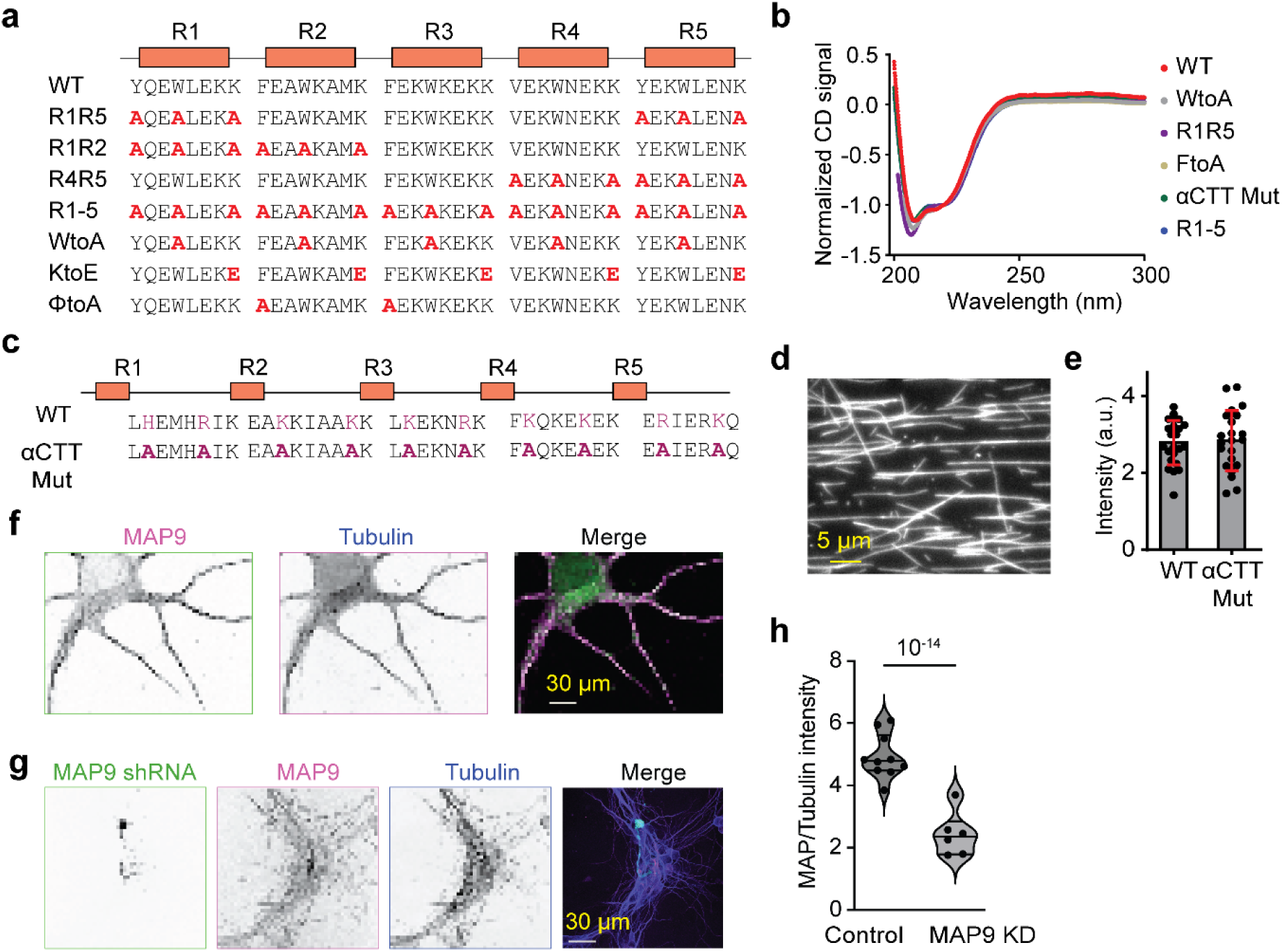
The analysis of MAP9 mutations in vitro and MAP9 knockdown in neurites. **a**, The schematic shows the mutations in MAP9 to disrupt the pairwise interactions between MAP9 and β-tubulin. The R index indicates that all three conserved residues were mutated to alanine in the specified pseudo-repeats. **b,** Circular dichroism (CD) spectra of WT and mutant MAP9-MTBD constructs fused to GFP. Spectra were normalized to the CD signal at 220 nm to compare the shape, regardless of protein concentration. Overall, the mutant constructs exhibit a similar CD spectrum shape to WT, indicating that the mutations do not substantially alter the overall secondary structure of the protein. **c,** Schematic of the αCTT mutant to disrupt potential interactions between MAP9 and the CTT of α-tubulin. **d,** Representative image of GFP-αCTT binding to MTs in 100 mM salt. **e,** Fluorescent intensity measurements reveal that αCTT and WT MAP9-MTBD decorate MTs with similar affinity (n = 20 MTs for each condition). **f,** Immunostaining of primary cultured neurons with antibodies against MAP9 (Invitrogen PA5- 58145, magenta) and alpha-tubulin (Sigma Clone DM1A T9026, green). MAP9 staining appears weaker than that of tubulin in the soma, suggesting that MAP9 does not bind to soluble tubulin and instead decorates polymerized MTs. **g,** Additional example image of DIV4 neurons transfected with MAP9 shRNA (green) at DIV0 and counterstained with antibodies against alpha-tubulin (Sigma Clone DM1A T9026, blue) and MAP9 (Invitrogen PA5-58145, magenta). **h,** MAP9 to tubulin intensity ratio in control and MAP9 shRNA-transfected neurons (n = 12 and 8 neurons, respectively, from 2 independent trials). p-values were calculated from Welch’s two-tailed t-test.

**Extended Data Fig. 5.**
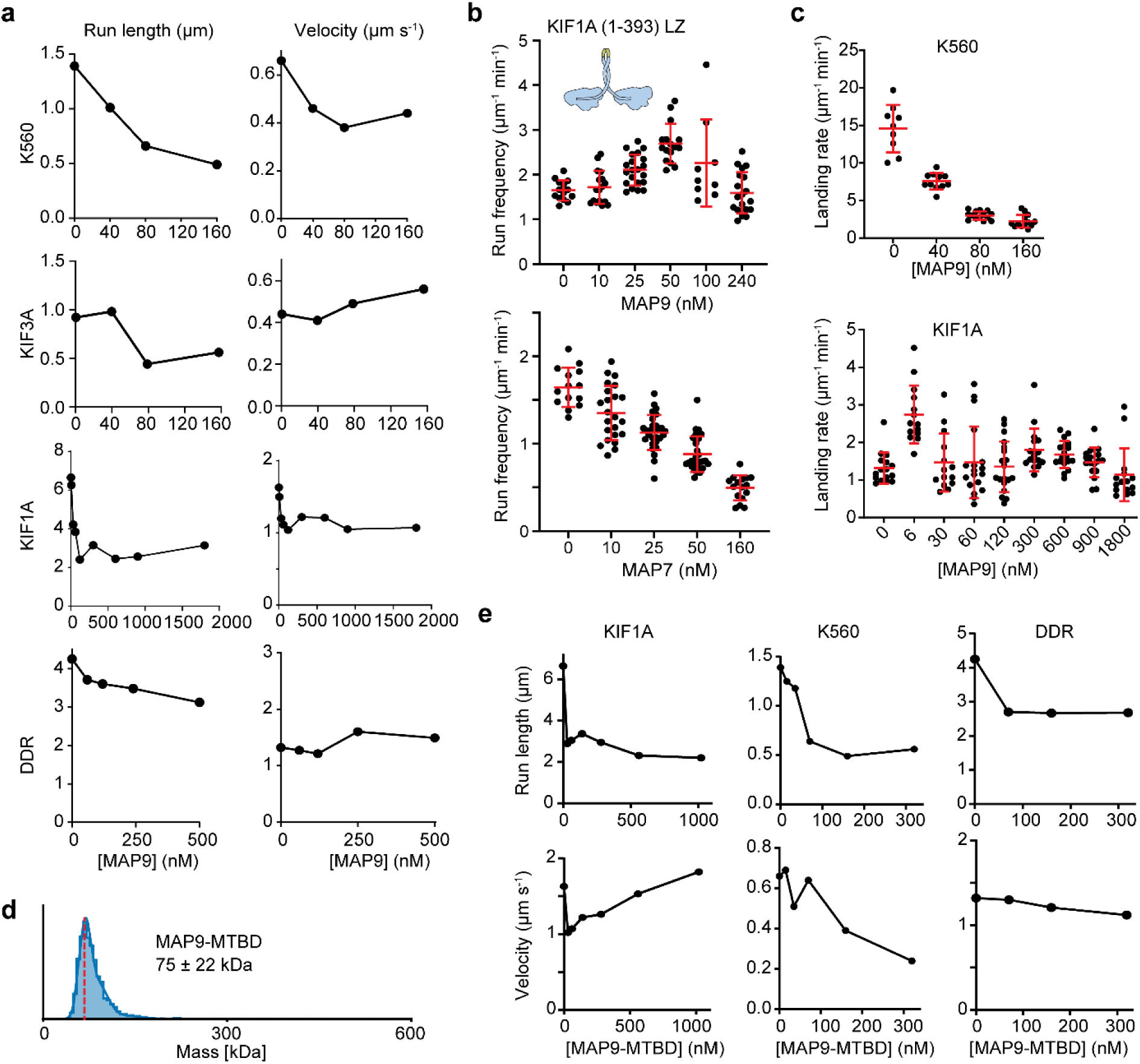
The analysis of in vitro motility of motor proteins on MAP-decorated MTs. **a**, The average velocity and run length of MT motors under different MAP9 concentrations. From left to right, n = 805, 580, 181, 105, motors for K560, 508, 156, 177, 74 motors for KIF3A, 384, 316, 379, 383, 173, 254, 373, 400, 130 motors for KIF1A, and 547, 527, 334, 226, 297 motors for DDR. **b,** (Top) Schematic of tail-truncated artificial dimer of KIF1A (LZ: leucine zipper). (Bottom) Run frequency of KIF1A (1-393)-LZ under different concentrations of MAP7 (from left to right, n = 14, 23, 28, 29, 16) and MAP9 (from left to right, n = 14, 17, 21, 16, 9, 19). The centerline and whiskers represent the mean and SD, respectively. **c,** The MT landing rate of K560 and KIF1A under different MAP9 concentrations. From left to right, n = 778, 2084, 1603, 415 motors for K560, and 1252, 697, 467, 1587, 725, 1010, 1324, 764, and 868 motors for KIF1A. **d,** Mass photometry of purified MAP9-MTBD-GFP (mean ± s.d.). The red dashed line represents the expected monomer mass. **e,** The velocity and run length of motors under different MAP9-MTBD concentrations. From left to right, n = 384, 290, 319, 446, 639, 502, 443 motors for KIF1A; 805, 705, 645, 298, 390, 339 motors for KIF5B; and 547, 518, 502, 408 motors for DDR.

**Extended Data Fig. 6.**
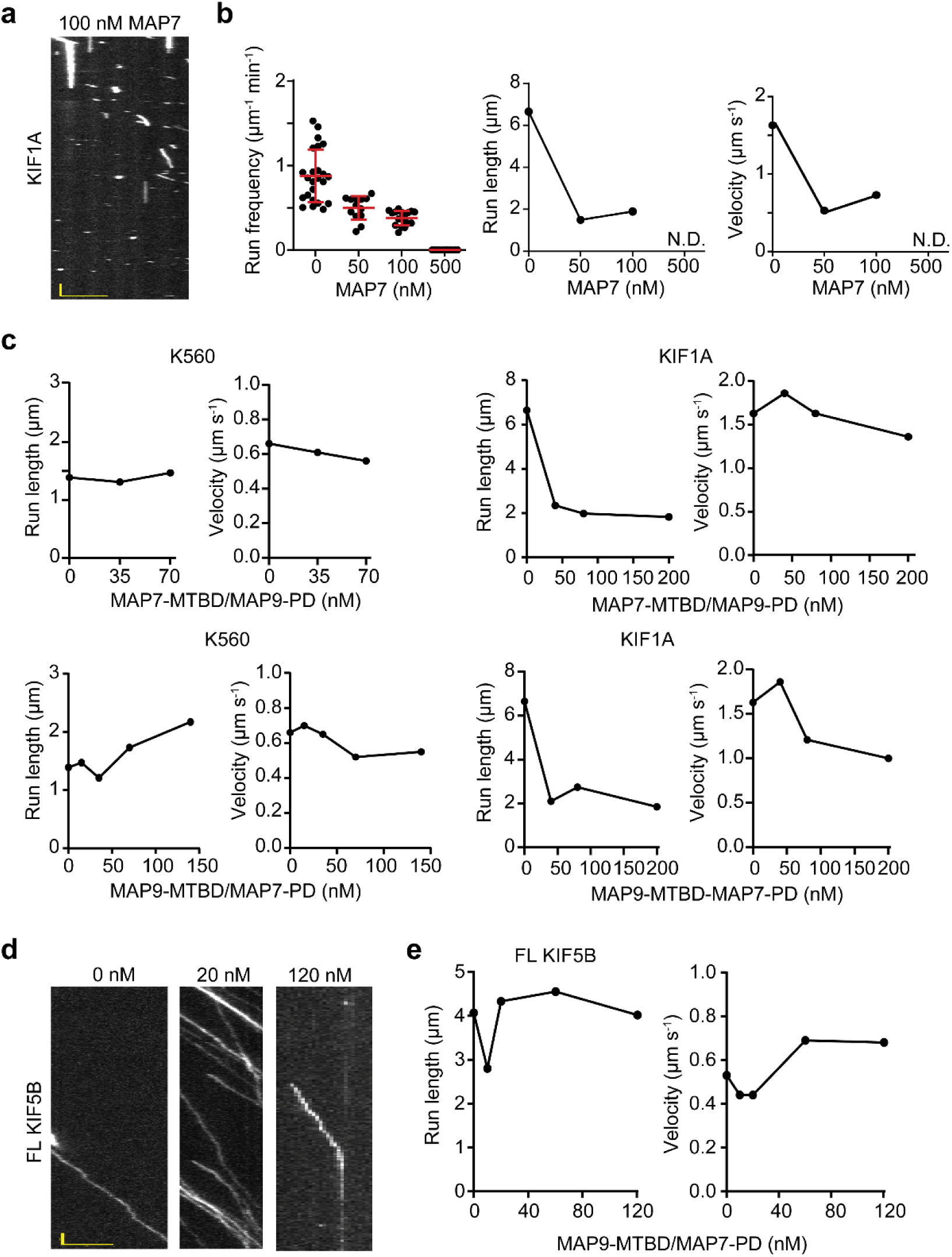
Regulation of KIF1A and KIF5B motility by chimeric MAPs of MAP7 and MAP9. **a**, MAP7 inhibits KIF1A. Kymograph of KIF1A motility in the presence of 100 nM MAP7. **b,** Run frequency, velocity, and run length of KIF1A in the presence of various amounts of MAP7. From left to right, n = 24, 13, 18, and 11 MTs for run frequency and 384, 96, and 142 motors for run length and velocity measurements. **c,** Run length and velocity of KIF5B and KIF1A under different concentrations of chimeric MAPs. From left to right, n = 805, 473, 224 motors for KIF5B and MAP7-MTBD/MAP9-PD, 805, 812, 409, 254, 226 motors for KIF5B and MAP9- MTBD/MAP7-PD, 384, 178, 152, 130 motors for KIF1A and MAP7-MTBD/MAP9-PD, and 384, 541, 537, 326 motors for KIF1A and MAP9-MTBD/MAP7-PD. **d,** Kymographs of FL KIF5B in the presence and absence of MAP9-MTBD/MAP7-PD chimera. **e,** The run frequency, run length, and velocity of FL KIF5B under different MAP9-MTBD/MAP7-PD concentrations. From left to right, n = 15, 28, 15, 21, 24 MTs for run frequency and 28, 393, 180, 76, and 76 motors for the run length and velocity measurements. In a and d, the scale bar is 5 µm on the x-axis and 5 s on the y-axis.

**Extended Data Fig. 7.**
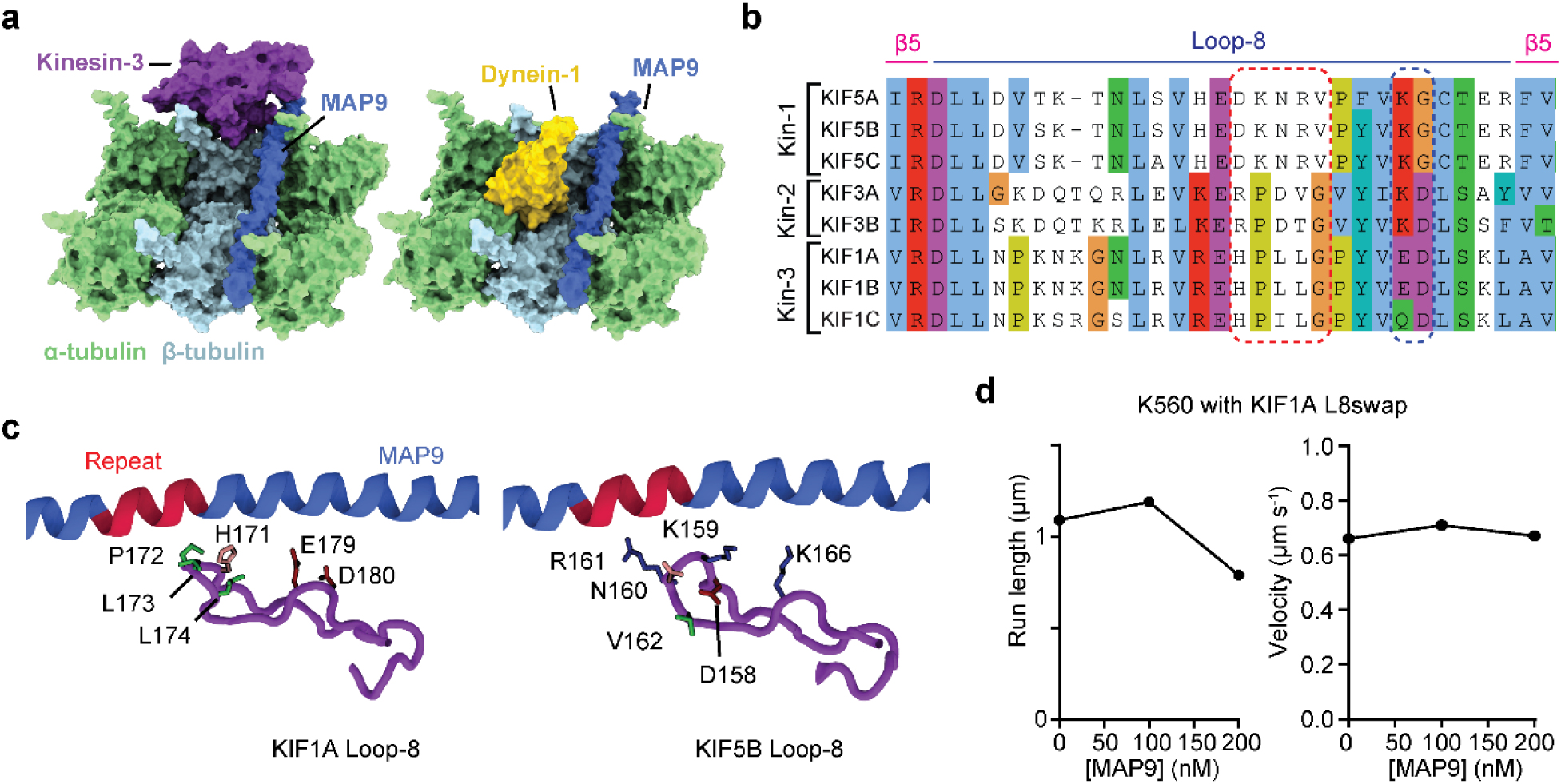
Loop-8 of the kinesin motor domain does not directly overlap with MAP9 on the MT. **a**, Alignment of the loop-8 region of transport kinesins from humans. MAP9 does not sterically overlap with kinesin or dynein on the MT. The cryo-EM structure of MAP9 was superimposed onto those of the kinesin-3 motor domain (PDB ID: 8UTO^39^) and dynein MTBD (PDB ID: 6RZB^60^), all bound to the MT, demonstrating that MAP9 occupies a distinct location without an obvious overlap with the footprint of these motors on the MT. **b,** The sequences are oriented from the N-terminus to the C-terminus, and members of the kinesin motor family were grouped based on their kinesin class. The sequences were aligned using the Clustal O algorithm in Jalview and colored based on the Clustal algorithm. **c,** Comparison of the atomic models of loop- 8 from kinesin-3 (this work) and kinesin-1 (PDB ID:3J8Y)^61^. Hydrophobic, negative, and positive side chains are highlighted in green, red, and blue, respectively. The sequence highlighted with a red dashed rectangle in a is most closely positioned to and interacts with the pseudorepeats of the MAP9 helix, and this sequence in kinesin-1 was swapped with that of KIF1A to generate the K560 L8swap mutant. The sequence highlighted with a blue dashed rectangle in b forms electrostatic interactions with the MAP9 helix. **d,** The run length and velocity of KIF5B L8swap under different concentrations of MAP9. From left to right, n = 631, 662, 420 motors.

**Extended Data Fig. 8.**
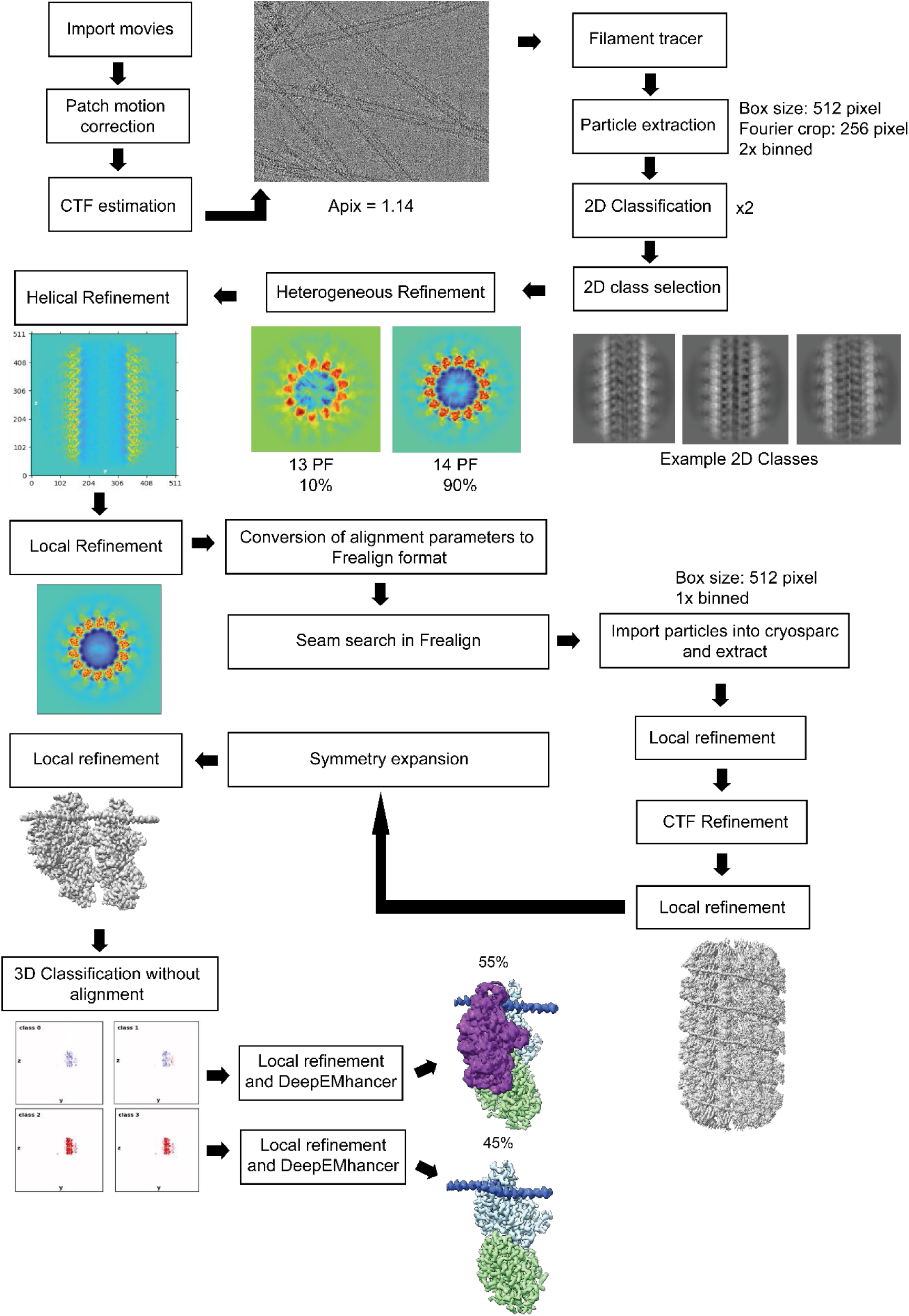
Cryo-EM data processing pipeline used to determine the MAP9- KIF1A-MT structure. The symmetry expansion was performed for two tubulin dimers laterally adjacent to each other.

**Extended Data Fig. 9.**
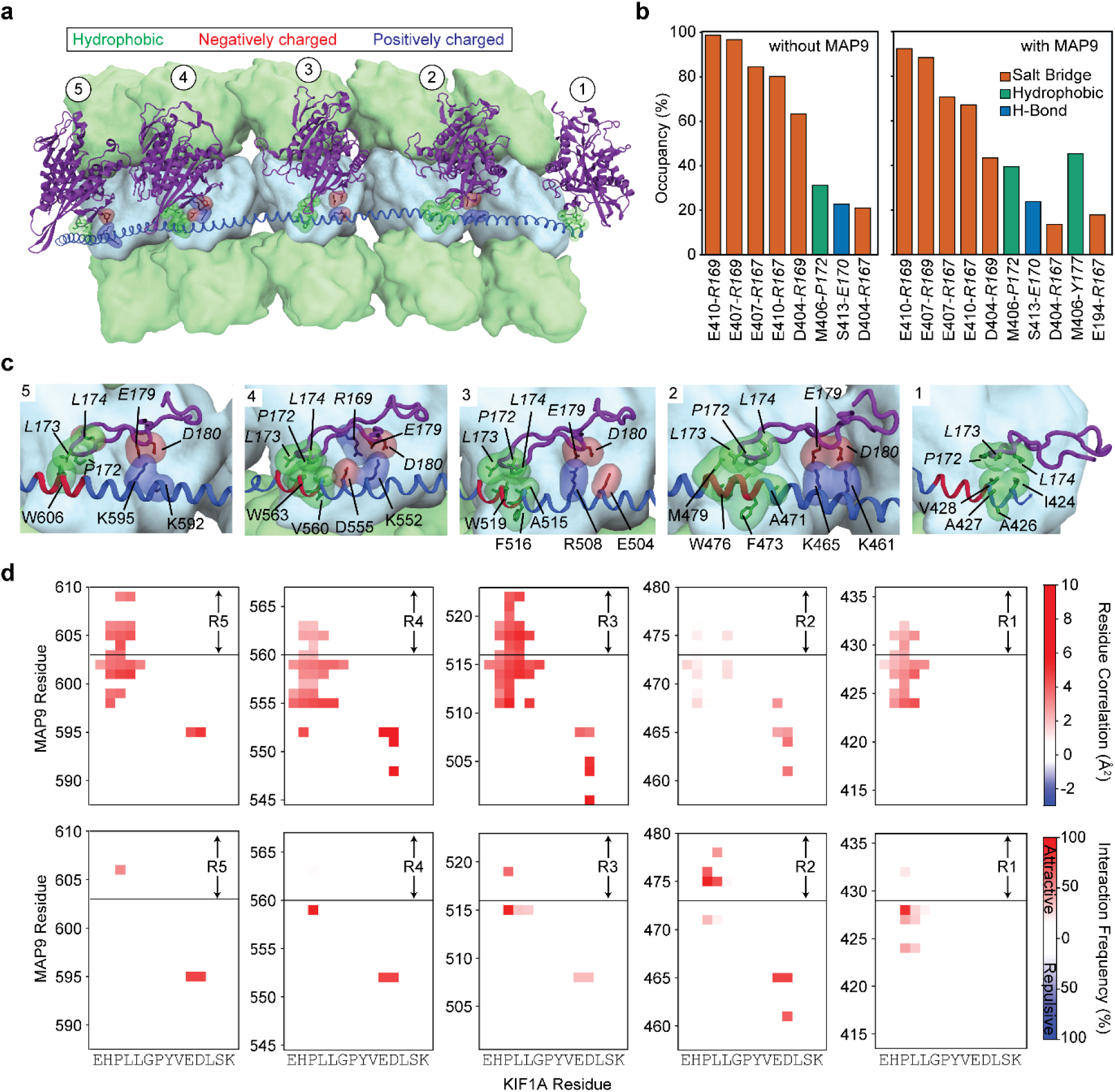
Interactions between MAP9 and kinesin-3 loop 8 from MD simulations. **a**, A representative snapshot from the MD simulation system containing five kinesin-3 motor domains (purple, cartoon representation), five tubulin lattice segments (surface representation; α-tubulin and β-tubulin in light green and light blue, respectively), and MAP9 (royal blue, cartoon representation), simulated in the presence of explicit solvent. **b,** The frequency of pairwise interactions between β-tubulin (regular font) and kinesin-3 loop-8 (italic) in the presence and absence of MAP9. **c,** Close-up structural views of each MAP9 repeat (R1–R5) highlighting the hydrophobic cores and electrostatic interactions formed between MAP9 amino acids (regular font) and kinesin-3 loop-8 (italic). **d,** (Top) Correlations between kinesin-3 and MAP9 amino acids throughout the MD simulations. (Bottom) Interaction frequency heat maps reveal the amino acid pairs responsible for these correlations. The magnitudes of attractive and repulsive correlations and interactions are color-coded.

**Extended Data Fig. 10.**
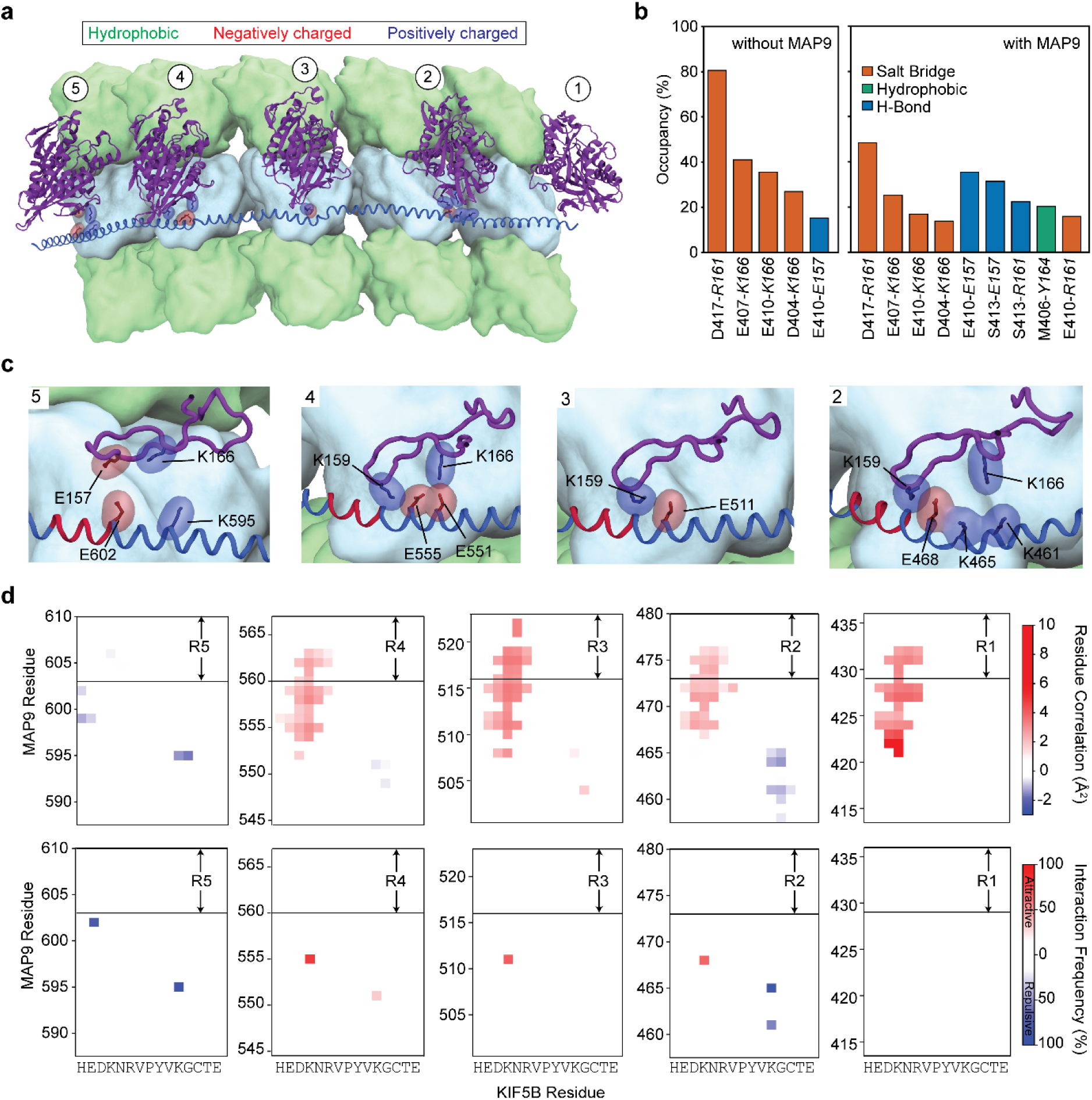
Interactions between MAP9 and kinesin-1 loop 8 from MD simulations. **a**, A representative snapshot from the MD simulation system containing five kinesin-1 motor domains (purple, cartoon representation), five tubulin lattice segments (surface representation; α-tubulin and β-tubulin in light green and light blue, respectively), and MAP9 (royal blue, cartoon representation), simulated in the presence of explicit solvent. **b,** The frequency of pairwise interactions between β-tubulin (regular font) and kinesin-1 loop-8 (italic) in the presence and absence of MAP9. **c,** Close-up structural views of each MAP9 repeat (R1–R5) highlighting the electrostatic interactions formed between MAP9 amino acids (regular font) and kinesin-1 loop-8 (italic). **d,** (Top) Correlations between kinesin-1 and MAP9 amino acids throughout the MD simulations. (Bottom) Interaction frequency heat maps reveal the amino acid pairs responsible for these correlations. The magnitudes of attractive and repulsive correlations and interactions are color-coded.

