## Supplementary Information for "Structure and mechanism of microtubule stabilization and motor regulation by MAP9"

### Supplementary Figures

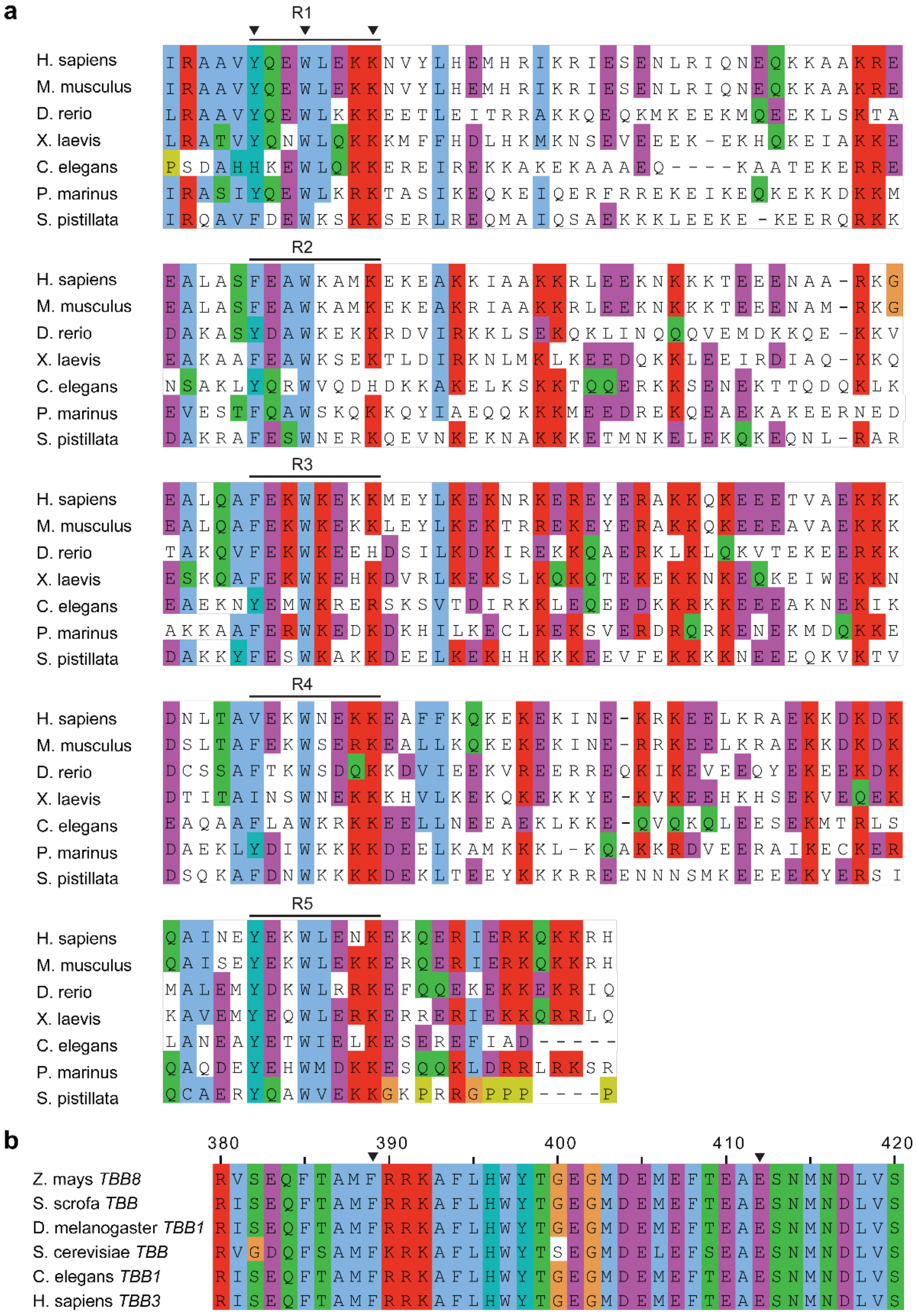


**Supplementary Fig. 1 | Sequence conservation of the MAP9-MT interface. a,** The sequence alignment of MAP9 MTBD across species reveals the conservation of its MT-binding repeats. The alignment covers the MTBD of human MAP9 (residues 424 - 624). **b,** The sequence alignment of β-tubulin across species reveals the conservation of its residues interacting with the MAP9 helix (arrowheads). The alignment covers residues 380 – 420 of human β3 tubulin. The sequences were aligned and colored using the Clustal O algorithm in Jalview.

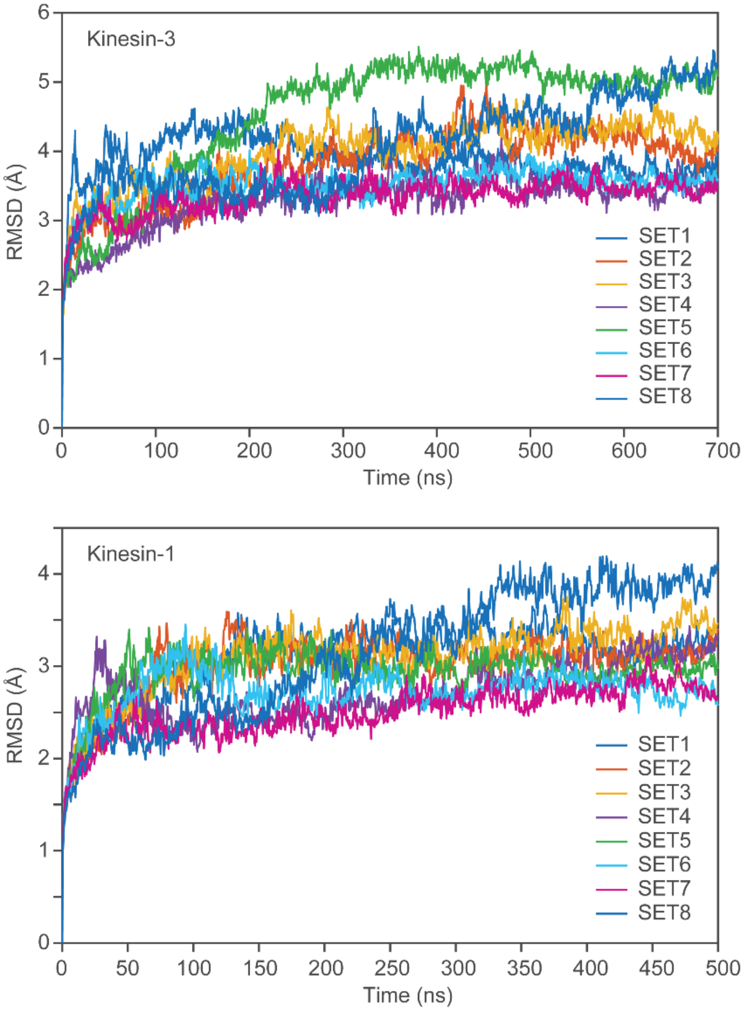


**Supplementary Fig. 2 | RMSD stability of kinesin–MAP9–MT complexes.** RMSD time traces for each independent MD trajectory of the kinesin-3–MAP9–MT complex and the kinesin-1–MAP9–MT complex relative to the starting conformation. Each colored trace represents one independent simulation. After the initial relaxation period, the RMSD values reach stable plateau regions, indicating that the complexes remain structurally stable during the production simulations without major conformational rearrangement or disruption of the MAP9–MT or kinesin–MAP9–MT interfaces.

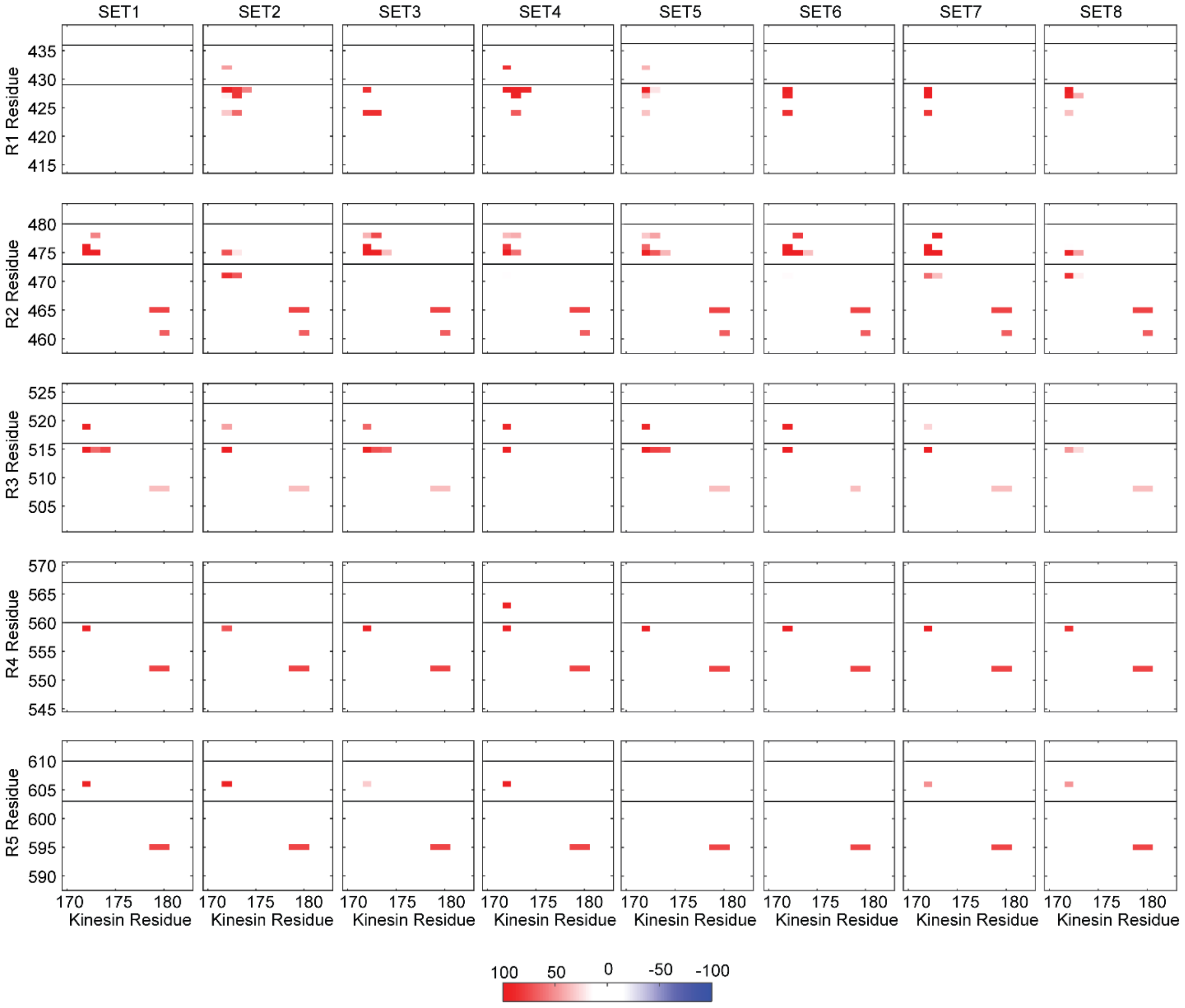


**Supplementary Fig. 3 | Trajectory-resolved MAP9–kinesin-3 interaction maps.** Interaction-frequency maps calculated separately for each independent MD trajectory of the kinesin-3–MAP9–MT system. The x-axis shows kinesin-3 loop-8 residues, and the y-axis shows residues of R1- R5 pseudorepeats of MAP9. Attractive/favorable interactions are shown in red, and repulsive/unfavorable interactions are shown in blue; color intensity indicates interaction frequency. The recurrence of similar interaction patterns across independent trajectories supports that the reported MAP9–kinesin-3 contacts are not derived from selected snapshots or a single simulation.

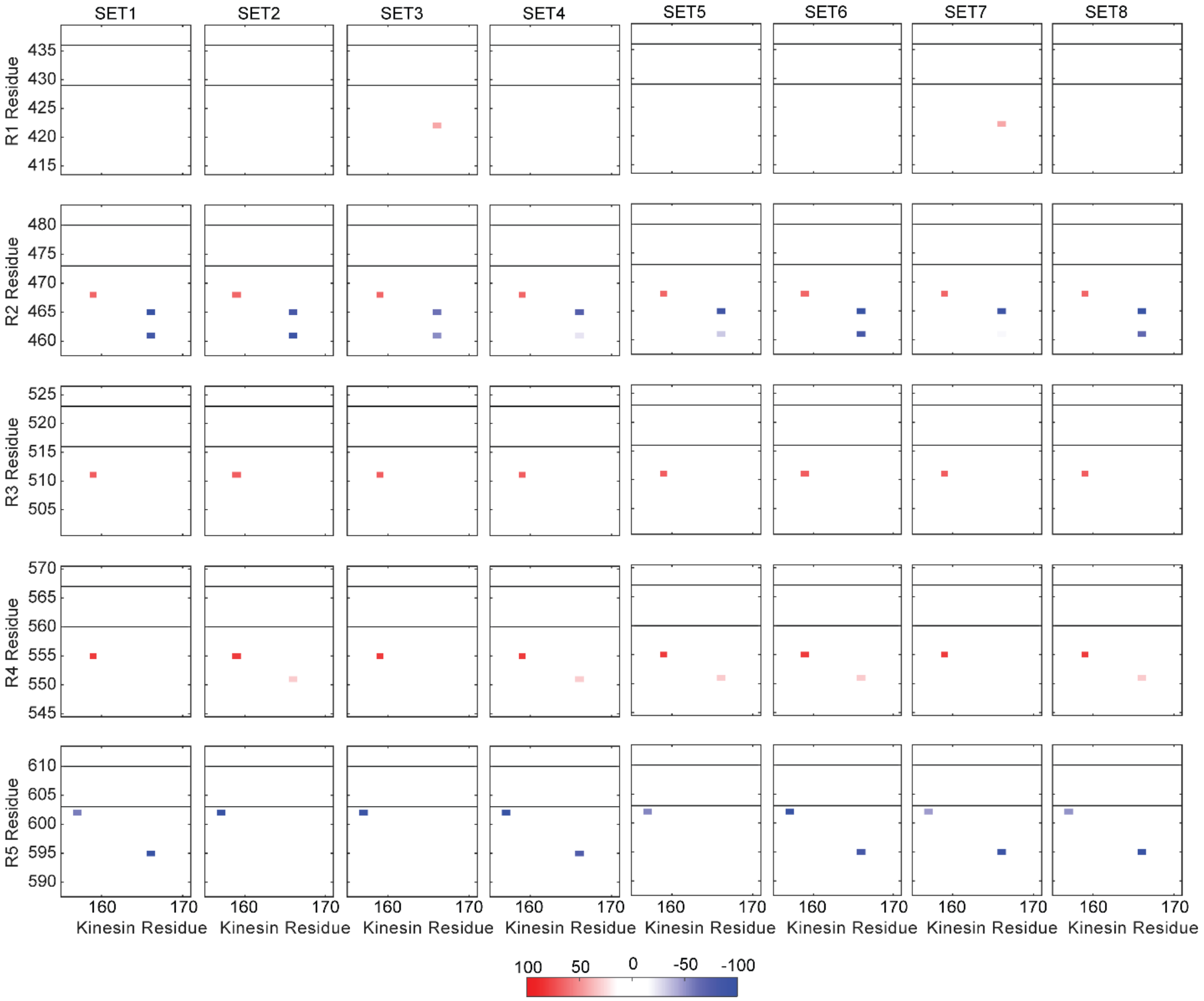


**Supplementary Fig. 4 | Trajectory-resolved MAP9–kinesin-1 interaction maps.** Interaction-frequency maps calculated separately for each independent MD trajectory of the kinesin-1–MAP9–MT system. The x-axis shows kinesin-1 loop-8 residues, and the y-axis shows residues of R1- R5 pseudorepeats of MAP9. Attractive/favorable interactions are shown in red, and repulsive/unfavorable interactions are shown in blue; color intensity indicates interaction frequency. The recurrence of similar interaction patterns across independent trajectories supports that the reported MAP9–kinesin-1 contacts are not derived from selected snapshots or a single simulation.

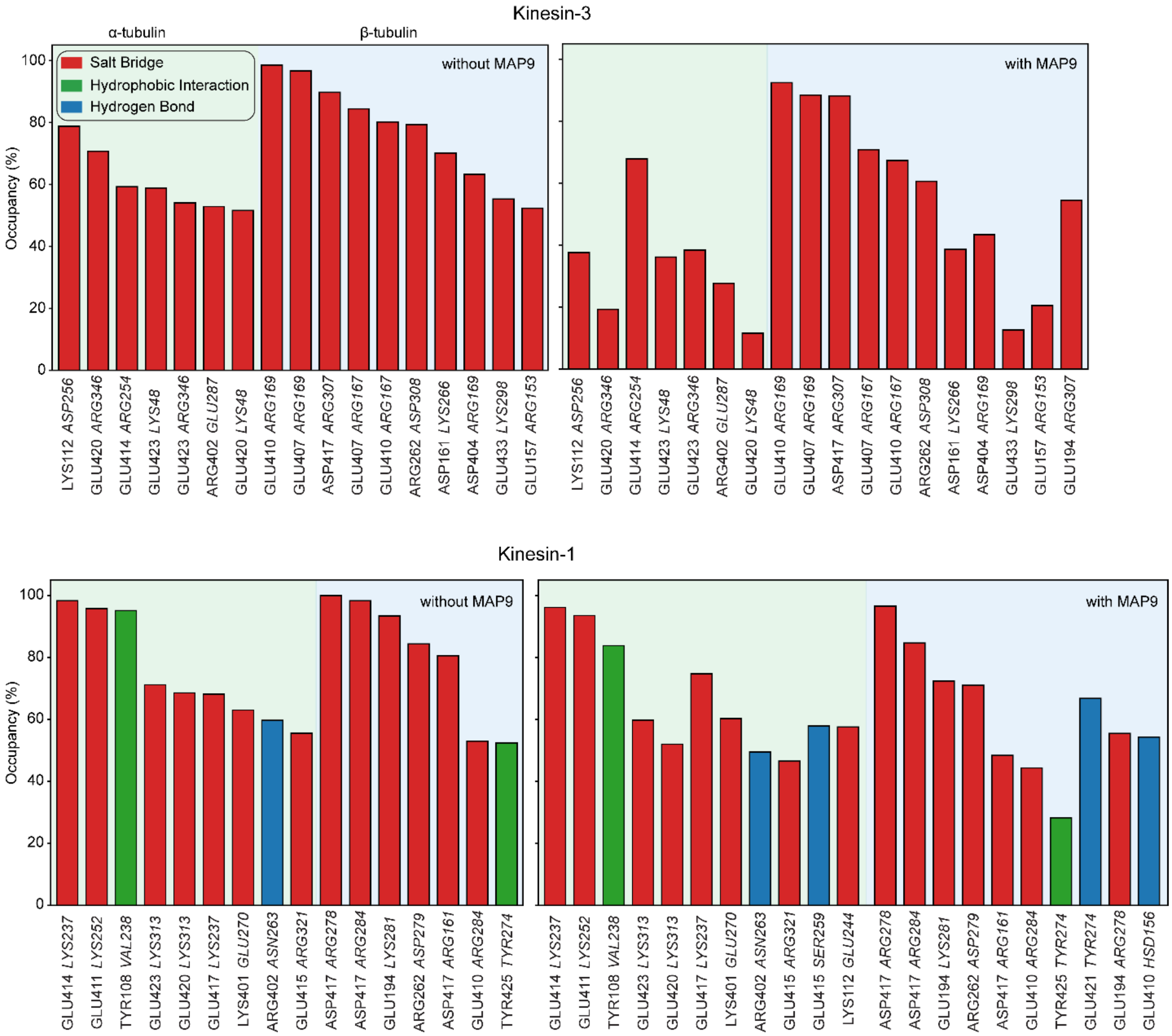


**Supplementary Fig. 5 | Kinesin-MT interactions in the absence and presence of MAP9.** MD simulations reveal residue-level interaction occupancies between kinesin-3 (above) or kinesin-1 (below) with the MT in the presence or absence of MAP9. Interactions are grouped according to the tubulin subunit (α- and β-tubulin) and classified as salt bridges (red), hydrophobic interactions (green), or hydrogen bonds (blue). Interacting kinesin and tubulin residues are shown on the x-axis using italic and regular fonts, respectively. Comparison with the MAP9-containing systems showed that MAP9 largely preserved the kinesin-1–MT interface, with a redistribution of salt-bridge and hydrogen-bond interactions, whereas it reduced kinesin-3 salt-bridge occupancy at the α-tubulin interface while preserving interactions with β-tubulin.

### Supplementary Tables

| Name | Description | Source | Expression | Figures |
|---|---|---|---|---|
| K560-GFP-HaloTag E215C CLM | Human KIF5B (1-560)$^{E215C}$ ::GFP::HaloTag (Cys-light) | [15] | E. coli | 4, 5, EDF5, EDF6 |
| FL KIF5B | Human KIF5B (1-963)::GFP:: SNAP | [15] | Sf9 | 5, EDF6 |
| FL MAP7 | Human ybbR::MAP7 (1-749) | [15] | Sf9 | EDF6 |
| FL MAP9 | GFP::Human MAP9::strepII | [16] | E. coli | 1, 3, 4, 5, 6, EDF5 |
| KIF3A homodimer | Mouse KIF3A motor domain (1 – 359) fused to the coiled-coil of *Drosophila* KHC (345 - 559) | [56] | E. coli | 3, EDF8 |
| FL KIF1A | Human KIF1A::yBBR::strepII | This study | Sf9 | 4, 5, EDF5, EDF6, |
| KIF1A-LZ | Human KIF1A (1-393): leucine zipper: yBBR- strepII | This study | E.coli | EDF5 |
| MAP9-MTBD | Human MAP9 (302-647)::GFP::strepII | This study | E.coli | 2, 5, EDF4, EDF5, EDF6 |
| MAP9-MTBD R1R2 | Human MAP9 (302-647)$^{Y429A, W432A, F473A, W476A, K480A}$ ::GFP::strepII | This study | E.coli | 2, EDF4 |
| MAP9-MTBD R1R5 | Human MAP9 (302-647)$^{Y429A, W432A, K436A, Y603A, W606A, K610A}$ :: GFP::strepII | This study | E.coli | 2, EDF4 |
| MAP9-MTBD WtoA | Human MAP9 (302-647)$^{W432A, W476A, W563A, W519A,W606A}$ | This study | E.coli | 2, EDF4 |
| MAP9-MTBD FtoA | Human MAP9 (302-647)$^{F473A, F516A}$::GFP::strepII | This study | E.coli | 2, EDF4 |
| MAP9-MTBD KtoE | Human MAP9 (302-647)$^{K436E, K480E, K523E, K567E, K610E}$::GFP::strepII | This study | E.coli | 2, EDF4 |
| MAP9-R4R5 | Human MAP9 (302-647)$^{V560A, W563A, K567A, Y603A, W606A, K610A}$ ::GFP::strepII | This study | E.coli | 2, EDF4 |
| K560 L8 swap | Human KIF5B (1-560), (DKNRV 158-162 replaced with HPLLG) ::mScarlet::yBBR::strepII | This study | E.coli | 6, EDF7 |
| MAP9-MTBD/MAP7-PD | StrepII:GFP::Human MAP9 (302-647)::Human MAP7 (307-749) | This study | E.coli | 5, EDF6 |
| MAP7-MTBD/MAP9-PD | StrepII:GFP::Human MAP7 (60-316)::Human MAP9 (1-301) | This study | E.coli | 5, EDF6 |
| KIF1A motor domain, monomeric | mTurquoise2::Human KIF1A (1-393)::strepII | This study | E.coli | 6 |
| SNAPf-Dyn-Phi | SNAPf-DYNH1C1$^{K1610/R1567E}$-IC2C-LIC2-Robl1-Tctex1-LC8 | [57] | Sf9 | 4, 5, EDF5 |
| BicDR1-SNAP | Mouse BicDR1 (1-577)::SNAP | [15] | Sf9 | 4, 5, EDF5 |

**Supplementary Table 1 | The list of constructs used (EDF: Extended Data Figure).**

| EMDB ID | MAP9 (R1-R2)-MT<br>EMD-74518 | MAP9 (R2-R3)-MT<br>----- | MAP9-KIF1A (1-393)-MT<br>EMD-74517 |
|---|---|---|---|
| **PDB ID** | 9ZP6 | 9ZP7 | 9ZP5 |
| **Data collection** | Arctica | Arctica | Arctica |
| **Camera** | K3 | K3 | K3 |
| Voltage | 200 kV | 200 kV | 200 kV |
| Electron exposure | 50 $e^-/Å^2$ | 50 $e^-/Å^2$ | 50 $e^-/Å^2$ |
| Magnification | 36000x | 36000x | 36000x |
| Pixel size | 1.14 Å/px | 1.14 Å/px | 1.14 Å/px |
| Defocus range | -0.8 to -2 | -0.8 to -2 | -0.8 to -2 |
| Total micrographs | 2,425 | 2,425 | 2,303 |
| Initial particle images (before symmetry expansion) | 463,970 | 463,970 | 354,241 |
| Final particle images (after symmetry expansion) | 4,639,700 | 4,639,700 | 3,542,410 |
| Map resolution | 3Å | 3Å | 2.9Å |
| FSC threshold | 0.143 | 0.143 | 0.143 |
| Sharpening B-factor | -80 | -80 | -60 |
| **Model refinement** | Phenix | Phenix | Phenix |
| Model Resolution | 3.2Å | 3.2Å | 3.3Å |
| FSC threshold | 0.143 | 0.143 | 0.143 |
| **Model composition** | | | |
| Non-hydrogen atoms | 14,571 atoms | 14,768 atoms | 9,854 atoms |
| Protein residues | 1820 | 1845 | 1231 |
| Ligands | 6 | 6 | 5 |
| **Validation** | | | |
| Clashscore | 5.72 | 5.72 | 7.12 |
| MolProbity score | 1.47 | 1.70 | 1.83 |
| Rotamers outliers (%) | 1.09 | 1.91 | 2.45 |
| **Ramachandran plot** | | | |
| Favored | 97.35 | 97.00 | 97.21 |
| Allowed | 2.65 | 3.00 | 2.71 |
| Outliers | 0 | 0 | 0.08 |

**Supplementary Table 2 | Cryo-EM dataset and model-building statistics.**

**a**

| MD ID | CTT Conformation | Constraints | Kinesin Presence | Duration |
|---|---|---|---|---|
| a-b | α-helix | α- and β- Tubulins | No Kinesins | 125 ns |
| c-d | α-helix | α- and β- Tubulins, MAP9 terminals | No Kinesins | 125 ns |
| e-f | Unstructured | α- and β- Tubulins | No Kinesins | 125 ns |
| g-h | Unstructured | α- and β- Tubulins, MAP9 terminals | No Kinesins | 125 ns |
| i-j | α-helix | α- and β- Tubulins | Kinesin-3 | 700 ns |
| k-l | α-helix | α- and β- Tubulins, MAP9 terminals | Kinesin-3 | 700 ns |
| m-n | Unstructured | α- and β- Tubulins | Kinesin-3 | 700 ns |
| o-p | Unstructured | α- and β- Tubulins, MAP9 terminals | Kinesin-3 | 700 ns |
| q-r | α-helix | α- and β- Tubulins | Kinesin-1 | 500 ns |
| s-t | α-helix | α- and β- Tubulins, MAP9 terminals | Kinesin-1 | 500 ns |
| u-v | Unstructured | α- and β- Tubulins | Kinesin-1 | 500 ns |
| w-x | Unstructured | α- and β- Tubulins, MAP9 terminals | Kinesin-1 | 500 ns |
| y-z | Unstructured | α- and β- Tubulins | Kinesin-3 | 800 ns |
| aa-bb | Unstructured | α- and β- Tubulins | Kinesin-1 | 800 ns |

**b**

| | Residues |
|---|---|
| **MAP9** | A421, H624 |
| **α-Tubulin** | R2-V9, Q11-H28, F53-E55, H61-D69, P72-T80, L92-T94, L117-Q128, G131-F138, I209-L217, Y224-S241, T271-A273, V288-T292, L318-G321, P325-V328, G354-N356, R373-C376 |
| **β-Tubulin** | E3-A9, Q11-H28, I30-D31, T35-Y36, D41-Q43, V49-Y51, E53-A54, G57-E69, T72-P80, I84-R86, N89-Q94, L117-D118, V120-D128, G132-F133, L135-S138, A206-K216, D224-F242, A271-L273, E288-D295, F317-G319, E325-E328, A352-D355 |

**Supplementary Table 3 | Conformations and constraints used in MD simulations. a.** The list of MD simulations performed in this study. The CTTs were modeled either as α-helices or unstructured peptides. Constraints were introduced at the amino acid residues of tubulin to minimize the fluctuations of tubulin subunits independent of one another. Additional constraints were introduced at MAP9 terminals to ensure MAP9 remains bound to the MT. Two sets of simulations were performed in the presence and absence of kinesin-1 and -3 motor domains using each combination of constraints and CTT models. **b.** The list of amino acids from MAP9 termini, α-tubulin, and β-tubulin with constrained $C_\alpha$ atoms during MD simulations.

**Supplementary Video Legends**

**Supplementary Video 1. Interactions between MAP9 and tubulin observed in all-atom MD simulations.** All-atom MD simulations highlighting interactions between MAP9 (royal blue) and the tubulin dimer (α-tubulin in green, β-tubulin in blue). Simulations contain five tubulin lattice segments and MAP9 MTBD in the presence of explicit solvent (not shown). Protein backbones are depicted in cartoon representation. Interacting side chains are shown as licorice. Positively charged residues are colored blue, negatively charged residues are red, and hydrophobic residues are green. The MAP-tubulin interactions are magnified in separate panels for regions R1 through R5. The combined trajectory has a total duration of 1 μs and comprises eight separate MD simulation runs. The trajectory time (in ns) and the corresponding MD run number for each displayed conformation are indicated in the top-left and top-right corners, respectively.

**Supplementary Video 2. MAP9 promotes MT polymerization and suppresses catastrophe.** Single-color imaging of MT polymerization (red) from GMP-CPP seeds (not shown) immobilized to the glass surface through biotin-streptavidin linkage. The fluorescence background is due to the presence of unpolymerized tubulin in the flow chamber. Images were acquired at 5 s per frame, with a pixel size of 160 nm.

**Supplementary Video 3. Processive motility of FL KIF1A in the presence and absence of varying MAP9 concentrations**. MAP9 inhibits KIF1A at high concentrations. One-color imaging of LD655-labeled FL KIF1A on surface-immobilized MTs in the presence of 0, 15, and 600 nM MAP9. Images were acquired at 200 ms per frame, with a pixel size of 160 nm.

**Supplementary Video 4. Processive motility of DDR complexes in the presence of MAP9 and MAP9-MTBD**. One-color imaging of LD655-labeled DDR on surface-immobilized MTs without a MAP, with 320 nM MAP9-MTBD and 500 nM MAP9. FL MAP9 inhibits DDR motility. However, no significant inhibition is observed with MAP9-MTBD. Images were acquired at 200 ms exposure per frame.

**Supplementary Video 5. Processive motility of KIF5B (K560) complexes in the presence and absence of varying concentrations of MAP9.** One-color imaging of LD655-labeled K560 on surface-immobilized MTs with no MAP9, and with 40 and 80 nM MAP9, respectively. MAP9 inhibits the MT binding and motility of K560. Images were acquired at 200 ms exposure per frame.

**Supplementary Video 6. Processive motility of FL KIF1A in the presence and absence of varying MAP9-MTBD concentrations**. MAP9-MTBD does not inhibit FL KIF1A. One-color imaging of LD655-labeled FL KIF1A without MAP9-MTBD, and with 280 and 560 nM MAP9-MTBD. Images were acquired at 200 ms per frame.

**Supplementary Video 7. Interactions between MAP9 and kinesin-3 loop-8 from MD simulations.** All-atom MD simulations highlight interactions between kinesin-3 motor domains (purple, cartoon representation) and MAP9 (royal blue, cartoon representation). Simulations contain five kinesin-3 motor domains, MAP9, and five tubulin lattice segments (surface representation; α-tubulin and β-tubulin colored in light green and light blue, respectively), and explicit solvent (not shown). Close-up structural views of each MAP9 repeat (R1–R5) highlight the hydrophobic cores and the salt bridges formed between MAP9 residues and loop-8 of kinesin-3.

**Supplementary Video 8. Interactions between MAP9 and kinesin-1 loop-8 from MD simulations.** All-atom MD simulations highlight attractive as well as repulsive interactions between kinesin-1 motor domains (purple, cartoon representation) and MAP9 (royal blue, cartoon representation). Simulations comprise five kinesin-1 motor domains, MAP9, five tubulin lattice segments (surface representation; α-tubulin and β-tubulin in light green and light blue, respectively), and explicit solvent (not shown). Close-up structural views of each MAP9 repeat (R1–R5) highlight salt bridges and electrostatic repulsions between MAP9 residues and loop-8 of kinesin-1.
